# Mucin induces CRISPR-Cas defence in an opportunistic pathogen

**DOI:** 10.1101/2021.08.10.455787

**Authors:** Gabriel MF Almeida, Ville Hoikkala, Janne Ravantti, Noora Rantanen, Lotta-Riina Sundberg

## Abstract

Parasitism by bacteriophages has led to the evolution of a variety of defense mechanisms in their host bacteria. However, it is unclear what factors lead to specific defenses being deployed upon phage infection. To explore this question, we exposed the bacterial fish pathogen *Flavobacterium columnare* to its virulent phage V156 in the presence of a eukaryotic host signal (mucin). All tested conditions led to some level of innate immunity, but the presence of mucin led to a dramatic increase in CRISPR spacer acquisition, especially in low nutrient conditions where over 60% of colonies had obtained at least one new spacer. Additionally, we show that the presence of a competitor bacterium further increases CRISPR spacer acquisition in *F. columnare*. These results suggest that ecological factors are important in determining defense strategies against phages, and that the concentration of phages on metazoan surfaces may select for the diversification of bacterial immune systems.

## Introduction

One of the central themes in host-parasite interactions is to understand the diversity and ecology of host defense strategies^1-4^. Variation in defenses increase plasticity: one defense strategy may be useful in one setting but inefficient or costly in another^1^. However, the ecological conditions that lead to selective deployment of different immune strategies are still poorly understood. Defense mechanisms may be especially complex in systems where multiple trophic levels are involved^5^. For example, pathogenic bacteria that infect eukaryotic hosts are also parasitized by their viruses, bacteriophages (phages). This tripartite cross-domain relationship involves two layers of interactions that are influenced by host defenses: one between the bacterial host and the phage, and another between the metazoan host and the pathogenic bacterium. The interplay of these layers is still poorly understood^6^, and the continuum of evolutionary interactions extend to even those between the phage and the metazoan^7-12^.

Traditionally, phage-bacterium interactions are studied in simplified laboratory conditions. However, in real life, the interactions between pathogenic bacteria and phages often occur on the mucosal surfaces of vertebrate hosts. Phages are important members of mucosal microbiomes^13^, and some have evolved a symbiotic relationship with eukaryotes by interacting with mucin glycoproteins on the mucosal surfaces^8^. This phage-metazoan mutualism enhances phages’ probability of encounter with bacterial hosts, while providing the host external mucosal immunity against invading bacteria^7-9^. Bacterial invasion into mucosa thus leads to competition for space, resources, and more importantly, subjects bacteria for potential phage infections. Interaction with the mucosa directly increases the virulence of many bacterial pathogens^7,14-15^. Recently, virulence upregulation by mucin exposure was correlated to increased susceptibility to phage infections in *Flavobacterium columnare* and *Aeromonas sp*^7^. Increased phage susceptibility in the mucosal environment might also play a role in the recent descriptions of a *Clostridium difficile* phage activity being improved by eukaryotic cells^11^ and the mucin enhancement of an *Escherichia coli* phage^10^. How the mucosal environment influence bacterial resistance against phages is so far unknown. If defenses against phages incur tradeoffs in virulence for pathogenic bacterial species, conflicts may emerge. Thus, defense strategies that minimize virulence trade-offs during colonization may be favored.

Bacterial defense mechanisms against phage infections are numerous and have evolved towards almost all phases of phage life cycles^16^. Surface modification (SM) is an extracellular defense response, in which the mutation, downregulation or deletion of cell surface proteins prevents phages from attaching to the cell^16^. Within the cell, restriction modification and CRISPR-Cas systems interrogate invading phage genomes by distinguishing them from host nucleic acids and targeting them for destruction. CRISPR-Cas is an adaptive immune system that stores genetic memories of phage genomes (spacers) into a repeat-spacer array in a CRISPR locus. Upon a subsequent infection, the array is expressed to CRISPR-RNA (crRNA)^17^. The resulting repeat-spacer oligos guide endonucleases, such as Cas9, to the spacer-complementary sequence in the phage genome that is then cut by the endonuclease^18-19^.

Since phage receptor proteins often play an important role in bacterial nutrient intake, secretion, motility or virulence, disruptions in these proteins generally impose fitness costs^16,20-21^. It has been suggested that these tradeoffs are important for maintaining phage-bacterium coexistence^21^. Costs may be permanent or short-lived: constitutive defenses, such as SM, impose continuous costs, whereas inducible defenses, such as CRISPR-Cas, minimize their costs by being generally activated only under specific stimuli. The investment in inducible or constitutive defenses depends on their relative costs in a given environment^3^. For example, surface receptors required for virulence can have major fitness costs if they are mutated during host colonization^22^.

Despite the sophistication of CRISPR-Cas, less than half of all bacterial species carry CRISPR-Cas loci^23^, implying they may induce fitness costs or be useful only in limited circumstances. Ecological variables such as temperature and oxygen levels are indeed important factors that correlate with CRISPR-Cas occurrence^24^. Interestingly, type II CRISPR-Cas systems seem to be enriched in pathogenic bacteria^24^, and CRISPR-Cas has been suggested to regulate bacterial virulence in some bacterial species^25^, e.g. in *Francisella novicida*^26^ and *Campylobacter jejuni*^27-28^. However, as multiple defenses may be beneficial^1^, bacteria with CRISPR-Cas do not rely exclusively on this defense. The relative costs of CRISPR-Cas have been found to increase with phage concentration, so that above a certain threshold, these costs exceed those of SM, leading to a shift in defense strategy^3^. Furthermore, presence of competing bacteria can select for increased CRISPR-Cas based resistance due to an amplification in SM-based costs^29^ The factors that influence defense strategies are still largely unknown and they may have important effects on phage evolution (e.g. via genomic mutations or the evolution of anti-CRISPR proteins) and population size.

We hypothesized that the mucosal interface may be a tipping point between CRISPR-Cas and SM defenses. We investigated this idea using the opportunistic fish pathogen *Flavobacterium columnare* that causes columnaris disease in freshwater fish^30^. This bacterium has type II-C and VI-B CRISPR-Cas loci, both active in natural and laboratory environments^31-32^. Exposure to high concentration of phage in laboratory conditions elicits SM defense in *F. columnare*, causing loss of virulence and motility associated with colony morphotype change^33^. Low phage pressure and low nutrient level, on the other hand, can be used to trigger CRISPR-Cas spacer acquisition^32^. Exposure to primary catfish mucus has been shown to upregulate several virulence genes, as well as the CRISPR-Cas adaptation gene *cas2*^34^.

How simultaneous exposure to host signals (mucin), competing bacteria and phage affects the choice of phage defense strategy between CRISPR-Cas and SM has not been previously explored, despite the mucosal environment being a hotspot for microbial interactions. We studied how low-nutrient and simulated mucosal environments influence the ecology of phage resistance, in co-cultures of *F. columnare* strain B245 and its virulent phage V156. We also evaluated the effect of bacterial competition (with *Aeromonas sp*. bacteria) for phage resistance in the simulated mucosal condition. We show that while SM is a central resistance mechanism for bacteria, mucosal environment and bacterial competition significantly increases CRISPR spacer uptake. CRISPR-Cas activation by mucin exposure could allow the bacterial pathogen to remain virulent while evading predation by mucosal phages. These results provide crucial information on the ecological factors that shape immune defense strategies that are relevant for bacterial colonization of a vertebrate host.

## Materials and methods

### Phage and bacteria

*Flavobacterium columnare* strain B245 was isolated from the same fish farm in Central Finland in 2009 as its phage V156^35^. Conventional culturing of B245 was made using Shieh medium^36^ without glucose. V156 stocks were produced by harvesting confluent soft-agar layers from double-agar plates, adding 4ml of media, centrifuging (10.000 rpm, 10 minutes, Sorvall RC34 rotor) and filtering the supernatant through 0.22 µm filters. *Aeromonas sp*. B135 was isolated from a small natural brook in Central Finland (2008). *E. coli* DSM613 strain was obtained from DSMZ GmbH (Braunschweig, Germany).

### Long-term culturing conditions and sampling

The effect of four different nutritional conditions on phage resistance mechanisms were studied: autoclaved lake water alone, lake water supplemented with 0.1% purified porcine mucin, 0.1x Shieh medium alone and 0.1x Shieh media supplemented with 0.1% purified porcine mucin. The lake water was collected from Lake Jyväsjärvi (Jyväskylä, Finland) on February 13^th^ 2018 and autoclaved. The water was analyzed by Eurofins Scientific and contained N: 790 µg L^-1^, P: 18 µg L^-1^ and Fe: 540 µg L^-1^. Shieh media was diluted in ultrapure sterile water. Autoclaved 2% w:v solution of purified porcine mucin (Sigma-Aldrich, catalog no. M1778) was used as stock for preparing the simulated mucosal cultures.

The initial inoculum in each culture was 5×10^4^ colony forming units (cfu) of *F. columnare* B245 and 5×10^3^ plaque forming units (pfu) of phage V156 (multiplicity of infection of 0.1). Each condition was tested in triplicates, in a final volume of five milliliters, and incubated at 26 degrees under 120 rpm. As non-infected controls, one culture of each condition was made with only B245 without phage.

Every week after day zero, one milliliter of each culture was removed and replaced with one milliliter of the corresponding culturing media (autoclaved lake water or 0.1x Shieh, supplemented or not with 0.1% mucin). An overview of the experiment setup is shown in Figure 1.

**Figure 1.**
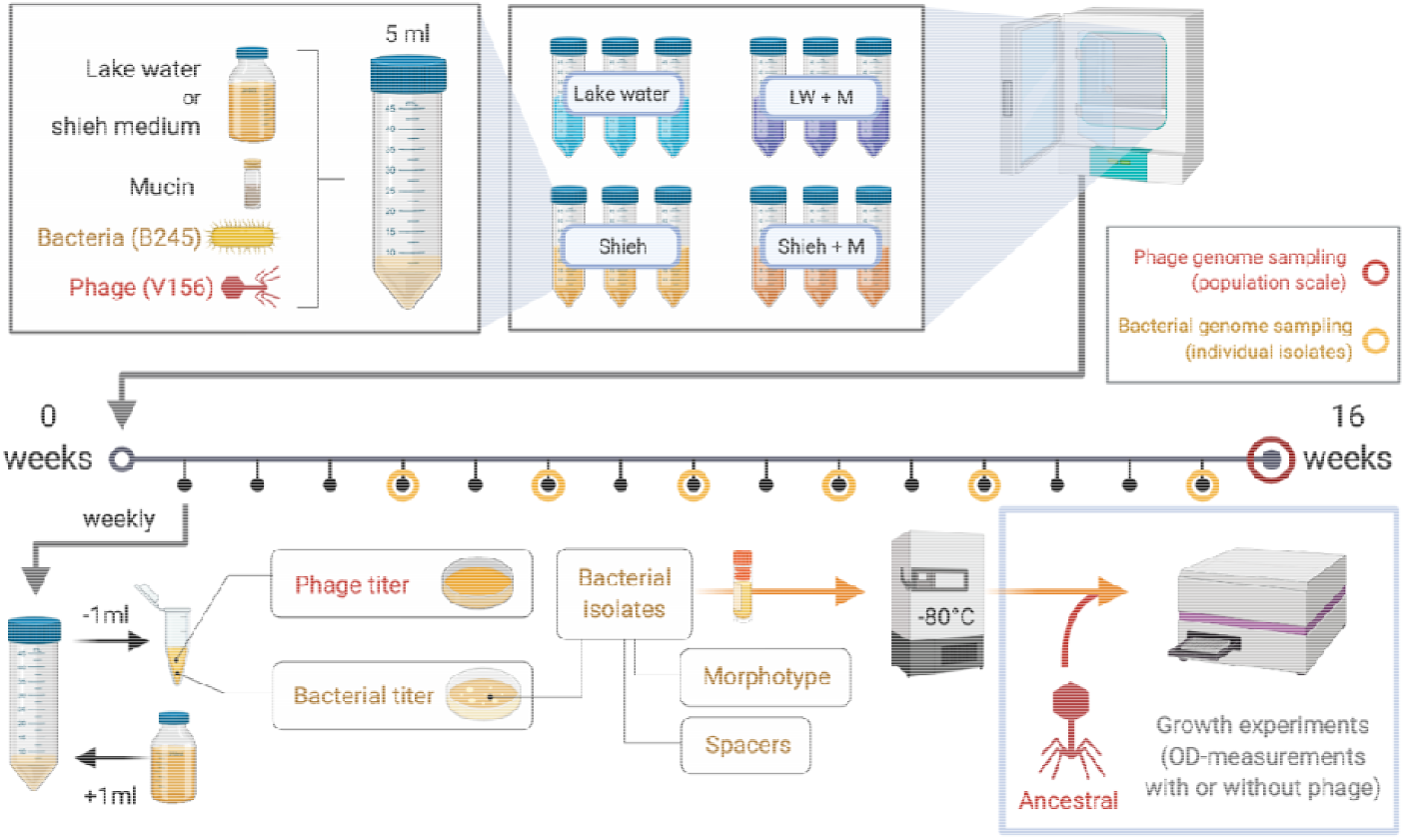
Overview of the experimental setup. The 16-week experiment (denoted by the horizontal line) contained four culturing conditions, which were sampled and restocked with fresh media weekly. Bacterial isolates were characterized by their morphotype and CRISPR spacer content and used later for growth tests with or without the ancestral phage. Phage genomes (population level) were sequenced at week 16. Genomes from representative bacterial isolates were sequenced throughout the experiment. Figure made in ©BioRender - biorender.com

### Bacteria and phage titrations

Immediately after sampling, each sample was serially diluted and plated on Shieh-agar plates for analysis of bacterial population size. Chloroform (10% v:v) was added to the remaining sample to kill bacterial cells. Serial dilutions of the chloroform-treated supernatants were used for titrating phages with the double-agar layer method^37^ using the ancestral B245 as host. Bacteria and phage plates were incubated at room temperature for three days, followed by the enumeration of bacterial colonies and viral plaques. In the first experiment phages were titrated every week during the experiment, while bacteria were titrated in weeks 1 to 8, 10, 12 and 15. The competition experiment was sampled at days 7, 14, 32 and 56 for phage and bacteria titers.

### Detecting the acquisition of new CRISPR spacers (both loci)

We aimed at collecting an equal number of Rough and Rhizoid colonies per sample whenever possible. After counting, random colonies were picked and transferred to 50 microliters of Shieh on 96 well plates. Two microliters of each resuspended colony were used as template for PCR reactions designed to detect the insertion of new spacers on both CRISPR loci. Reactions were made with DreamTaq polymerase (Thermo Fisher), in 20 μL reactions containing 0.5mM of DNTPs and 0.5 μM of each primer using previously published primers for strain B245^32^. Cycling conditions were 95 degrees for 3 minutes followed by 30 cycles of 95 degrees for 30 seconds, 60 degrees for 30 seconds, 72 degrees for one minute and a final extension step of 72 degrees for 15 minutes. PCR reactions were resolved in 2% agarose gels and the addition of spacers verified by the size of each amplicon.

### Bacterial growth characteristics

Representative bacterial isolates, considering colony morphology and CRISPR loci sizes, were chosen every week for further analysis. Following PCR, the remaining volume of resuspended colonies was transferred to 5mL of Shieh media. After overnight growth (120 rpm, 26 degrees) the individual colonies/isolates were frozen at -80 degrees for future use, and revived overnight for testing their immunity against the ancestral phage using Bioscreen C® (Growth curves Ltd, Helsinki, Finland). 1000 cfu mL^-1^ of each isolate were added to Bioscreen plates, in triplicates (200 microliters per well). Each isolate was tested in the presence and in the absence of the ancestral V156 phage (10^3^ pfu mL^-1^, MOI 1). Ancestral bacterial strain B245 was included on every plate as control, also in triplicates. Optical density measurements were made every 10 minutes for 4 days. Plates were kept without agitation at 27 degrees for the whole time. Minor differences between plates were accounted for by including the plate ID as a random effect in statistical models when appropriate. However, differences between plates was generally minimal (Supplementary Figure 1). Testing the growth of bacterial isolates was not made for the competition experiment (see below).

### Genomic DNA extraction and sequencing

Population-level phage DNA at week 16 and clonal DNA of selected bacterial isolates from different time points were sequenced with Illumina (BGI). For bacterial genomic DNA extraction, isolates were taken from the freezer and grown overnight. DNA of turbid cultures was extracted using the GeneJet Genomic DNA Purification Kit (Thermo Fisher). For phage DNA extraction, lysates from week 16 were used to infect B245. Confluent soft-agar bacterial lawns were collected, mixed with 4ml of Shieh media, centrifuged (10.000 rpm, 10 minutes, Sorvall RC34) and filtered. Phage precipitation was made with ZnCl2 followed by removal of host DNA with nucleases^38^. After Protease K treatment, the material was mixed with Guanidine:Ethanol (1 part 6M guanidine and 2 parts 99% ethanol, v.v.) and the extraction finished using the GeneJet Genomic DNA Purification Kit (Thermo Fisher). All samples were sequenced using 150PE BGISEQ platform at BGI Group. We were unable to obtain phage sequence data from the replicate b of the lake water with mucin condition due to technical problems.

#### Construction of the B245 reference genome

The ancestral B245 genome was assembled from Illumina reads using Spades 3.14.1 (--isolate mode)^39^. The resulting 487 contigs were combined to a single contig relying on a previously compiled complete *F. columnare* genome FCO-F2 (accession number CP051861) as reference using RagOO 1.1^40^. The genome was annotated for the purpose of mutational analysis using dFast 1.2.3^41^.

#### Mutation analysis

We used Breseq 0.35.1^42^ to analyze mutations occurring in the phage and bacterial genomes. Since the phage samples represented mixed phage populations, Breseq was run in --polymorphism-prediction mode. Bacterial samples were run in default mode. As references we used the ancestral B245 bacterial genome or the previously published V156 phage genome^31^. To reduce noise in phage mutation, we used a value of 0.05 for --polymorphism-bias-cutoff. During analysis of the phage genomes, we discovered a large number of identical polymorphisms in some samples. As it is very unlikely that such mutations occur independently, we discarded any identical mutations that co-occur in two or more samples as false positives. When investigating mutations in unknown putative phage genes, we used HHpred to predict protein function^43^.

### Competition experiment

A second experiment was made using only the lake water supplemented with 0.1% mucin condition (LW + M). The experimental setup was similar to the growth experiment described above, but besides adding 5×10^4^ cfu *F. columnare* B245 and 5×10^3^ pfu phage V156, we also included 5×10^4^ cfu of *Aeromonas sp*. B135 and of *E. coli* DSM613 as competitors. Cultures containing all three bacteria and V156, any combination of two bacteria and V156, and only *F. columnare* B245 host and V156 were made. Another control consisted of cultures containing only the V156 phage to follow its stability over time in the absence of its host. Each condition was tested in triplicates, in a final volume of five milliliters, and incubated at 25 degrees under 120 rpm. For this competition experiment we followed phage and bacterial titers, spacer acquisition in *F. columnare* and colony morphology.

### Statistics/Data analysis

All data analysis and statistic were done in R 3.5.3 using RStudio 1.1.463. Model comparisons and inclusion of random effects was aided by Akaike’s information criterion (AIC) comparison when applicable^44^. For details, see Supplementary Information.

## Results

### Presence of mucin stabilizes survival of both the bacterium and the phage during 16 weeks of co-culture

To avoid population bottlenecks, our sampling was based on weekly collecting 20% of the cultures and replacing with the same volume of fresh medium. Long term co-existence of both *F. columnare* B245 and its phage V156 was observed in all treatments. In lake water with (LW+M) or without mucin (LW) the phage titers remained similar until week 9, after which LW+M showed a significant decline in phage numbers compared to LW (LM, T_46_=-2.737, P = 0.0087) with roughly a ten-fold difference at week 16 (Figure 2A, Supplementary Figure 2A). Bacterial population densities in these treatments were opposite and more dramatic: bacterial numbers in LW+M were an average of 45-fold higher across all time points after an initial spike at week 1 (LM, T_77_=3.893, P < 0.001) (Figure 2B, Supplementary Figure 2B). Surprisingly, bacteria in the no-phage control of LW+M became extinct after week 10, while no extinction occurred in the phage-containing cultures.

**Figure 2.**
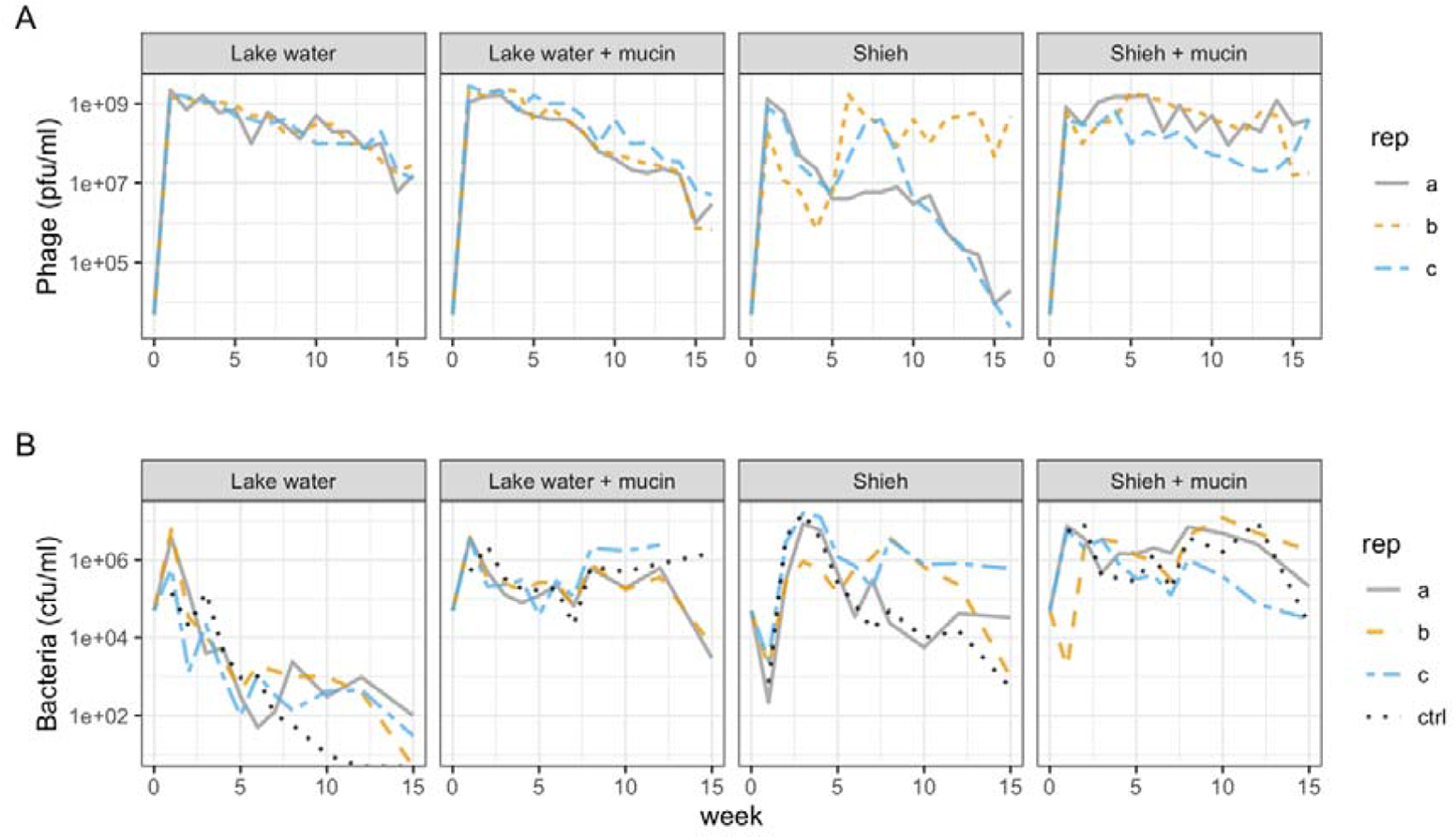
Phage V156 (A) and *F. columnare* B245 (B) titers over the 16-week experiment in the four treatments. Each line represents one of the three replicates in each treatment. The dotted lines in (B) are control cultures without phage.

Cultures in Shieh medium (without mucin) had large variation between replicates in both bacterial and phage titers. Despite similar bacterial titers at week one, differences between replicates grew to more than 100-fold towards the end of the experiment (Figure 2B). Phage titers declined in Shieh for the first 4-5 weeks, after which replicates *b* and *c* recovered while replicate *a* stabilized. Around week 10, phage titers declined sharply in replicates *a* and replicate *c*, while replicate *b* remained with high titer until the end (Figure 2A). Presence of mucin in Shieh decreased variation between replicates compared to Shieh alone, with higher phage and bacterial titers (Figure 2). No statistical analysis was performed on the Shieh cultures due to high divergence in replicates.

### Presence of mucin enhances spacer acquisition in type II-C and VI-B CRISPR-Cas loci

Culture conditions, especially the presence of mucin, had a significant impact in the acquisition of new CRISPR spacers (Figure 3A-B). The presence of mucin in lake water increased spacer acquisition 7.14-fold compared to plain lake water (GLMM, Z_3,68_ = -3.718, P=0.0037), and the efficiency of acquisition was 2.43-fold when compared to Shieh with mucin (GLMM, Z_3,68_ = -2.909, P<0.001) (Figure 3B). Shieh without mucin did not show any new spacers. The number of spacers in one locus in a single isolate was up to six spacers in LW+M, two in LW and up to three in Shieh with mucin (Figure 3B). The efficiency of spacer acquisition was roughly similar between the two CRISPR loci regardless of the treatment. Timewise, the maximum efficiency of acquisition was reached around week 3 in the LW+M treatment, with over 60% of LW+M colonies having acquired new spacers (Figure 3A). These data also show that spacer acquisition is not exclusive to the Rhizoid morphotype. In fact, in Shieh + M treatment, isolates with most spacers were Rough, indicating overlap of surface modification and spacer acquisition (Figure 3B). However, after a peak in isolates with CRISPR spacers in this treatment around week 7, newly acquired spacers disappeared on week 10. When only considering morphotype, LW had no Rhizoid colonies while other conditions had a minor bias towards them (Figure 3C).

**Figure 3.**
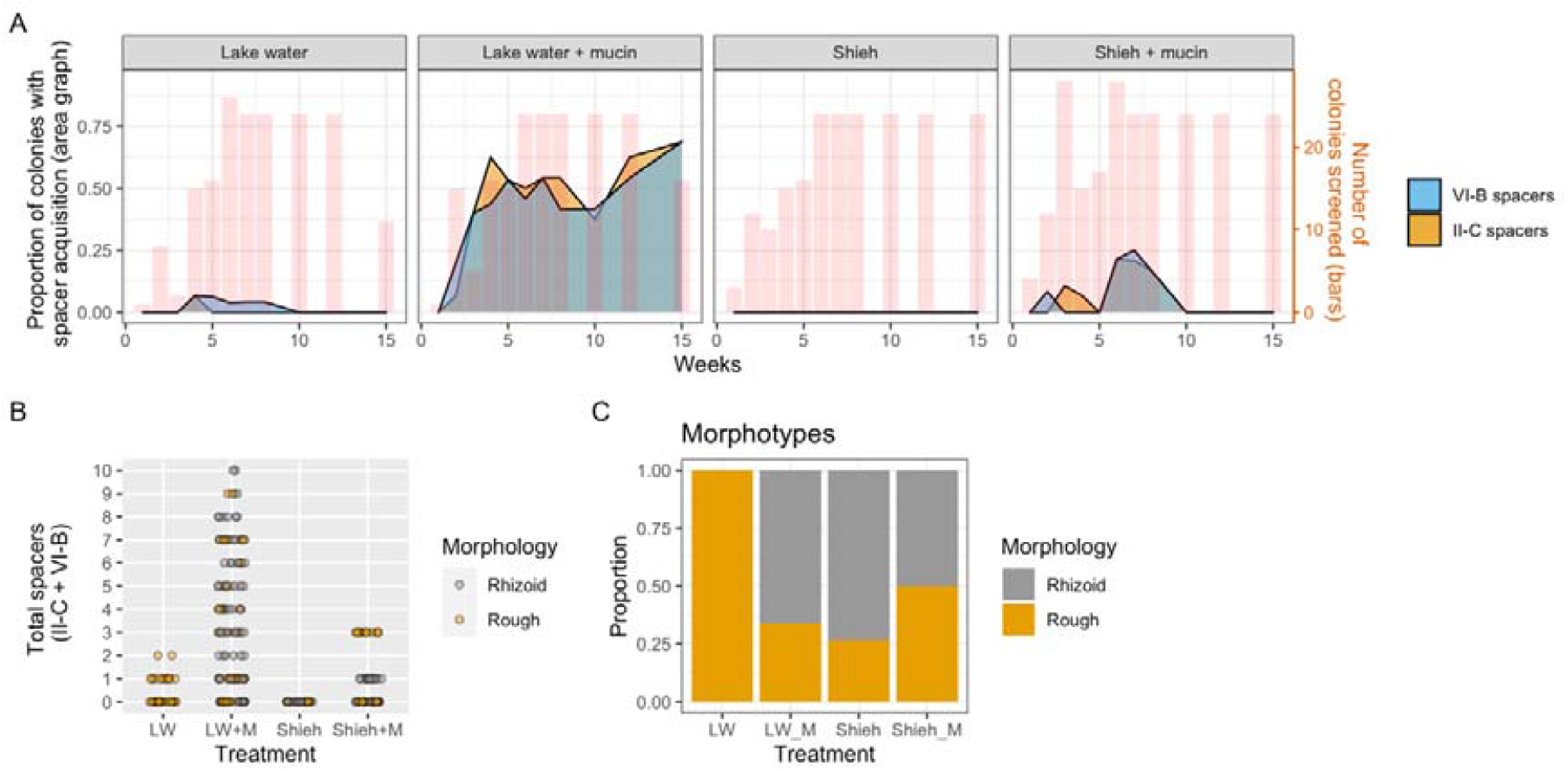
Dynamics of CRISPR spacers and colony morphotypes. A) CRISPR-Cas spacer acquisition over time in both loci. The area graphs (left Y-axis) represents the proportion of colonies in which the CRISPR-Cas array expanded by at least one new spacer. Values are means from the three replicates. The red line (right Y-axis) shows the total number of colonies that were screened at each time point to obtain these proportions. Missing red bars indicate no screening on that week – the respective proportional data is therefore an interpolation. B) The absolute number of spacers from individual isolates in different treatments. New spacers in both loci have been added together C) Morphotype distribution across the treatments (isolates from all time points pooled together).

### Co-culturing with phage leads to immunity

To detect the development of phage resistance and any associated costs during the 16-week co-culture experiment, we grew bacterial isolates obtained during the experiment in the presence or absence of the ancestral phage and compared the maximum OD (OD_MAX_) reached to that of ancestral B245. In the absence of phage, the OD_MAX_ of the four treatments (LW, LW+M, Shieh, Shieh+M) did not significantly differ from the ancestral B245 (Figure 4A). In the presence of phage, however, the OD_MAX_ of the ancestral B245 decreased from 0.41 to 0.19 (GLMM, Z_4,241_=-13.78, P < 0.001), while OD_MAX_ in the four treatments remained unchanged, indicating phage resistance (GLMM, Z_4,241_=-0.281 to -0.599, P > 0.3 in all) (Figure 4A). When compared with the ancestor in the presence of phage, the predicted maximum ODs were significantly higher in all treatments (GLMM, Z_4,241_=11.631 to 13.590), P < 0.001 in all) (Figure 4A). These results suggest that co-culturing *F. columnare* with phage caused phage resistance to evolve in all four treatments.

**Figure 4.**
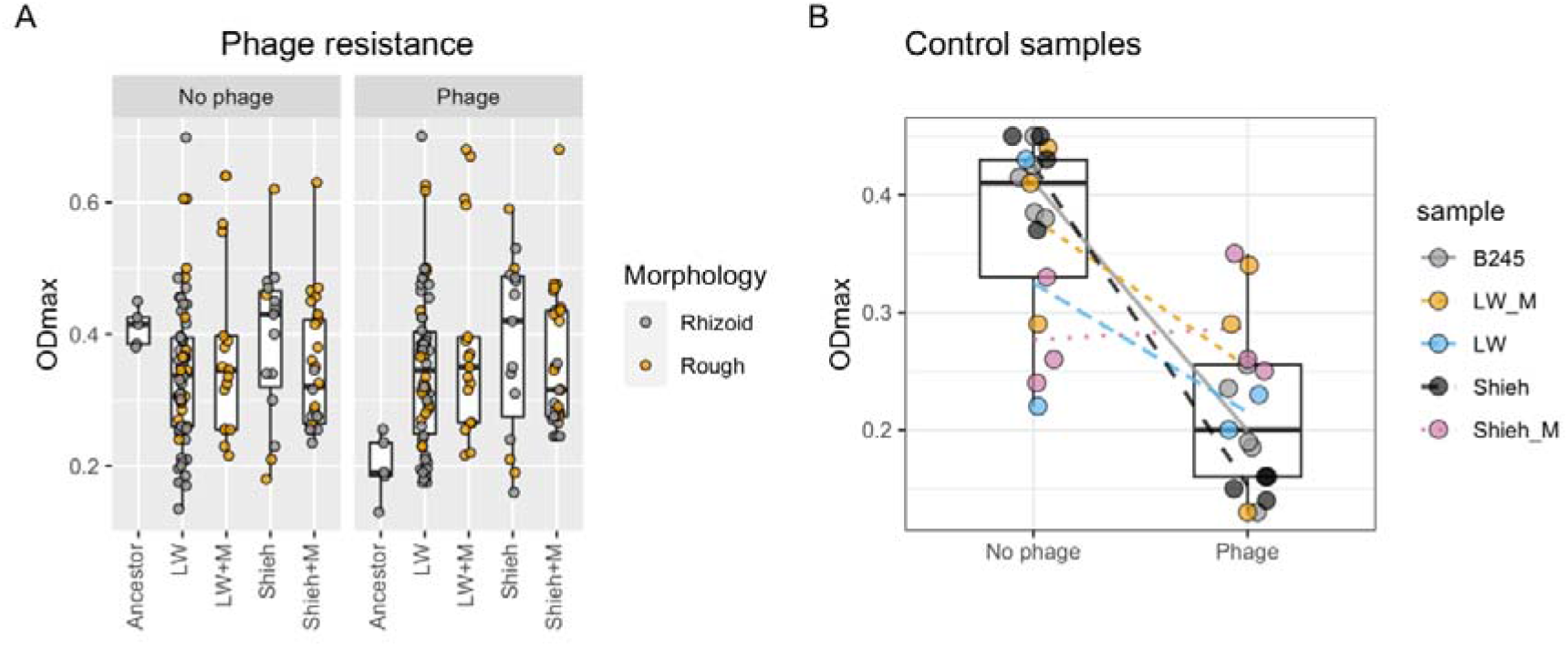
Effect of environment and colony morphology on bacterial growth. Each dot represents the mean of three replicate OD measurements of an isolate. A) Ancestral B245 and isolates originating from the four treatments in the presence or absence of phage. B) Control samples (grown without phage) from the same four treatments plus the ancestor in the presence or absence of phage. The box plots represent all treatments, including ancestor bacteria.

We also measured isolates from the control cultures from the 16-week experiment (same conditions but without phage). As expected, isolates from most treatments had lower OD_MAX_ in the presence of phage compared to the absence of phage (Figure 4B). The only exception was the Shieh + M isolate, which had a significantly lower OD_MAX_ in the absence of phage compared to the ancestral bacterium (OD 0.41) with a predicted OD of 0.276 (LM, T_2,24_=-3.053, P = 0.0055). In the presence of phage, however, this control had a predicted OD of 0.287, which is significantly higher than that of the ancestor’s 0.199 (LM, T_2,24=_3.565, P = 0.0016). These results suggest that prolonged incubation in Shieh + M in the absence of phage made the cells grow slower, but also made them phage-resistant, perhaps incidentally through adaptation to the nutrient-rich mucin environment.

### Fitness benefits of CRISPR adaptation depends on the environment

We next investigated how spacer acquisition in different morphotypes affected bacterial growth. This analysis was only done for the LW+M and Shieh+M treatments, which produced enough isolates with expanded CRISPR arrays for statistical analysis. In the absence of phage, Rhizoid and Rough isolates with native CRISPR arrays from the LW+M treatment reached predicted OD_MAX_ of 0.173 and 0.421, respectively (Figure 5A). This difference was statistically significant (GLMM, Z_1,50_=6.34, P < 0.001). However, acquisition of one or more spacers almost doubled the predicted OD_MAX_ of Rhizoid isolates from 0.173 to 0.337 (GLMM, Z_1,50_= 5.56, P < 0.001). The effect of new spacers was opposite for rough isolates, whose OD_MAX_ was reduced by 20% to a predicted 0.340 (GLMM, Z_1,50_=-5.326, P < 0.001). These effects were similar and significant also in the presence of phage (Figure 5). Sanger sequencing of selected bacterial isolates revealed that all sequenced spacers in II-C were targeting the phage (16/16), while roughly two thirds (19/28) were targeting the phage and the rest (9/28) targeting the bacterial genome in VI-B loci (Supplementary table 1).

**Figure 5.**
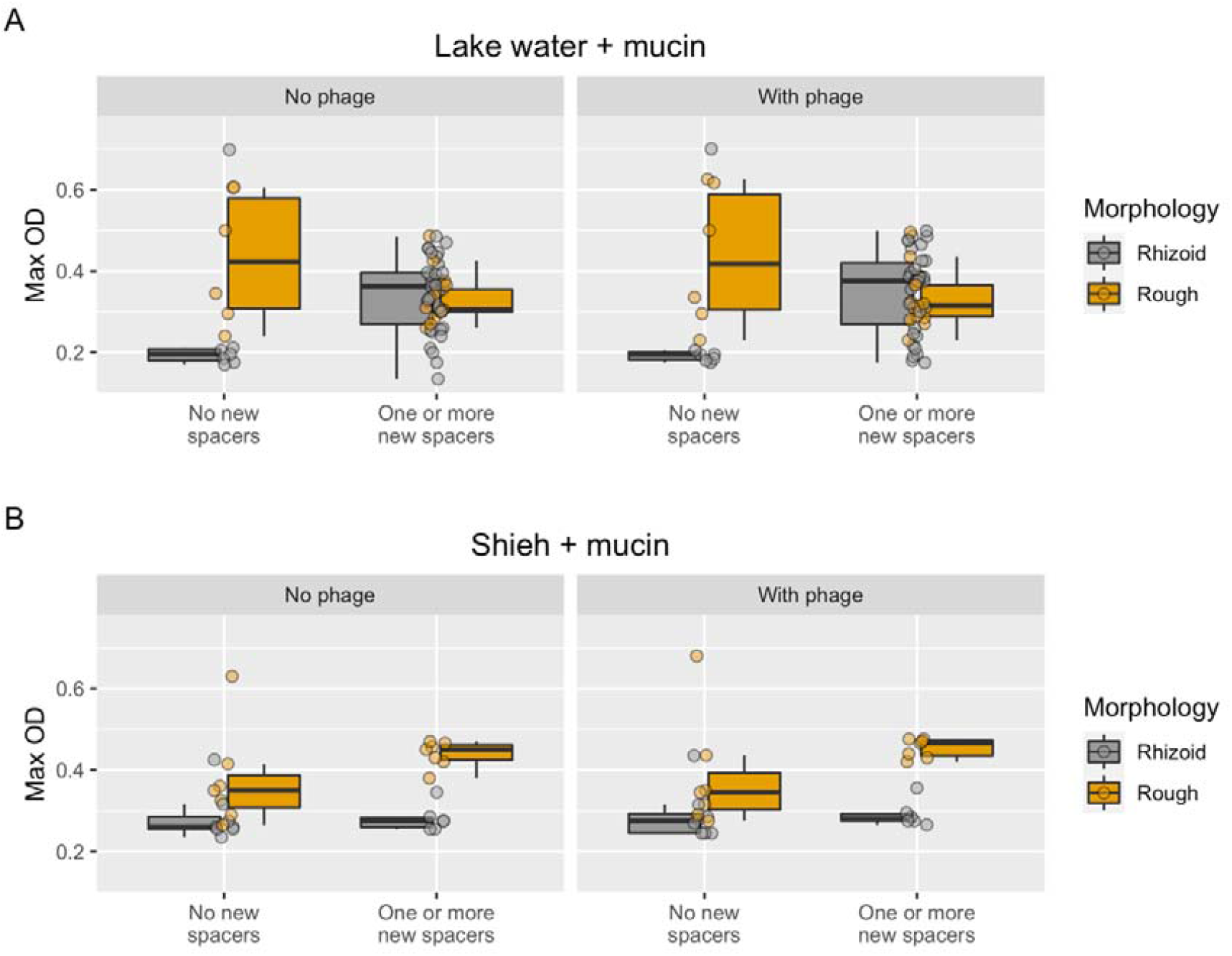
The effect of spacers and morphotype in LW+M and Shieh+M isolates on bacterial growth. Isolates are grouped into having no new spacers or having one or more new spacers regardless of the CRISPR-Cas locus.

In Shieh+M samples without phage, Rough isolates had a significantly higher OD_MAX_ than Rhizoid ones (0.400 vs 0.339, GLMM, Z_1,21_=2.240, P = 0.025) (Figure 5B). Surprisingly, new spacers increased OD_MAX_ of Rough isolates by 40% (GLMM, Z_1,21_=2.589, P = 0.01) but did not affect Rhizoid OD_MAX_. Again, results were similar when these isolates were subjected to phage (Figure 5B).

These results suggest that the conditions where CRISPR spacer acquisition first occurs influences bacterial growth in later conditions. CRISPR spacer acquisition has a positive impact on Rhizoid colonies and a negative impact on Rough colonies when the isolates originate from an environment that enhances spacer acquisition (LW+M, Figure 1A). However, when isolates originate from a less CRISPR-favouring environment (Shieh + M, Figure 1A), there is no or little benefit from the acquired spacers.

### Competition in lake water supplemented with mucin enhances spacer acquisition

After detecting that mucin in lake water caused an increase in CRISPR spacer acquisition, we performed a follow-up experiment testing the effect of competing bacterial species on *F. columnare* spacer acquisition in this condition. Initially we chose *Aeromonas sp*. which responds to mucin similarly as *F. columnare* by forming biofilm and by becoming more susceptible to phage infections, and *E. coli* DSM613 for apparently not being affected by mucin exposure^7^. However, the *E. coli* populations were either extinguished quickly (no colonies seen in the first samplings) or remained at very low levels (a few colonies seen at the last time point) during the experiment (Supplementary figure 4). This, allied to the fact that we could not measure which effect the initial input of *E. coli* had to mucin content in the cultures, led us to discard the *E. coli* -containing conditions from the final analysis. The presence of *Aeromonas sp*. significantly enhanced *F. columnare* spacer acquisition as measured by total number of spacers acquired across CRISPR loci (Figure 6). In the absence of *Aeromonas sp*., the expected total number of new spacers in *F. columnare* was 0.57 per colony, whereas in a co-culture each colony was expected to acquire 1.64 spacers (GLMM, Z=-3.381, P < 0.001). Interestingly, in this experiment the Rhizoid colony type bacteria acquired more spacers than the Rough types (Figure 6).

**Figure 6.**
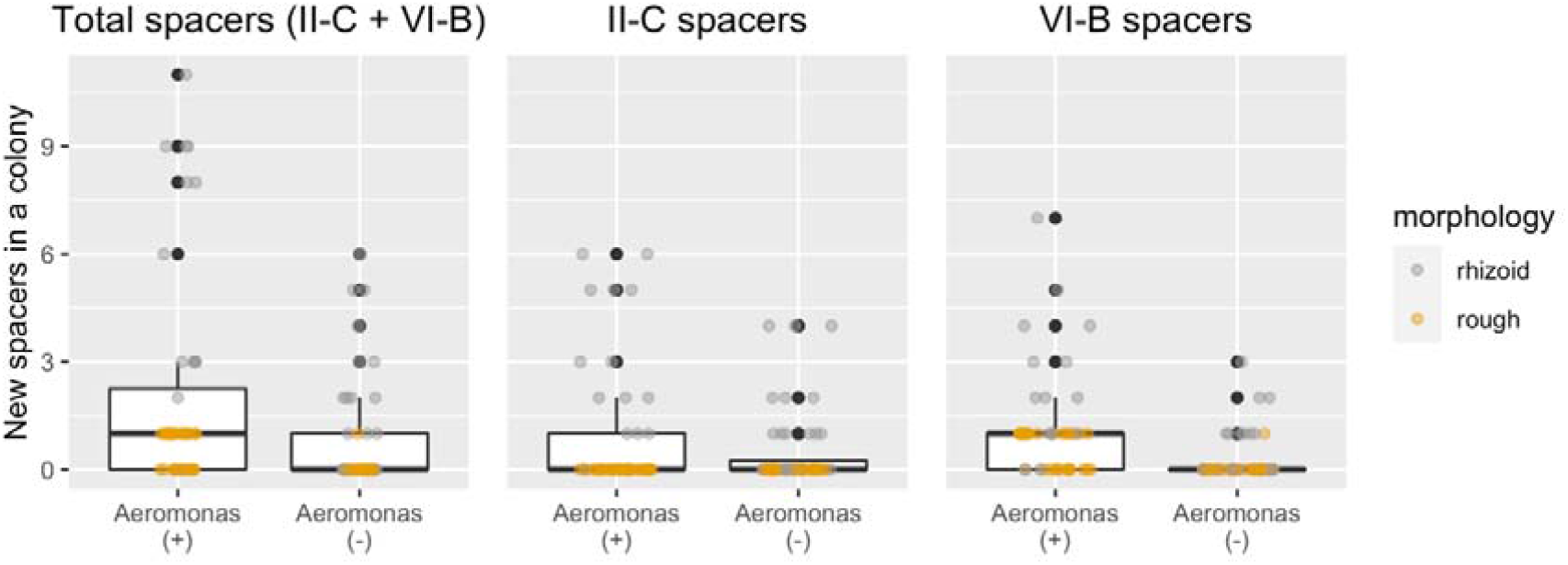
Effect of competitor on CRISPR-Cas spacer acquisition. The Y-axis shows the total number of spacers in a single *F. columnare* colony in the presence or absence of *Aeromonas*.

### Genome analysis

We sequenced phage and bacterial genomes to search for genetic variations resulting from coevolution. We also sequenced the control bacterial genomes (evolved without phage) to differentiate between mutations caused by interaction with phage and those that may arise from different culturing conditions.

Phage genomes were investigated on the population level: at the end of the 16-week experiment, phage samples of each culture (representing the variety of phages present in that culture) was used to infect the ancestral bacterium and the resulting lysate was deep sequenced. We found an abundance of shared mutations across multiple replicates. Due to the low likelihood of the same mutations occurring convergently across multiple samples, we discarded most mutations as false positives (however, all mutations are listed in Supplementary File 1). We were confidently able to recover individual mutations in only two cultures: Replicate *c* from Shieh + M treatment had a non-synonymous mutation (A_357_T) in a predicted phage baseplate protein and a nonsynonymous mutation (M_200_V) in a putative protein with no predicted function (Table 1). Replicate *b* from Shieh treatment had a non-synonymous mutation in an unknown protein near the aforementioned baseplate protein (V_300_I), as well as a synonymous mutation (K_10_K) in a predicted DNA helicase (Table 1). Common to both replicates is that they showed a drop in phage titer after the initial spike but recovered towards the end (Figure 2A). In fact, replicate *b* in Shieh treatment was the only replicate in its treatment group in which the phage titer remained high at the end of the experiment.

**Table 1.**
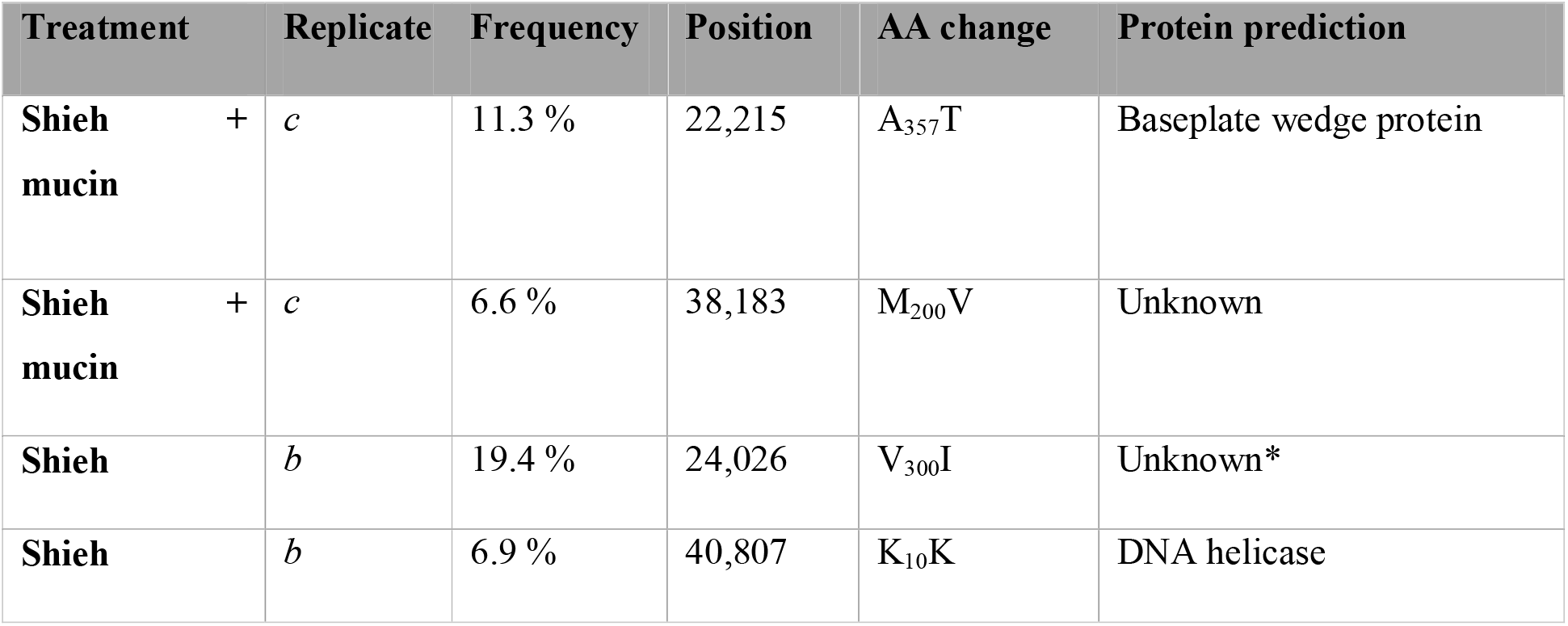
Mutations discovered in phage genomes on the population level. The frequency column shows the percentage of reads the mutation was found in. Position refers to the B245 genome. *while the function of this protein could not be predicted, its genomic neighborhood suggests a role as a phage structural protein.

| Treatment | Replicate | Frequency | Position | AA change | Protein prediction |
| --- | --- | --- | --- | --- | --- |
| Shieh + mucin | *c* | 11.3 % | 22,215 | A<sub>357</sub>T | Baseplate wedge protein |
| Shieh + mucin | *c* | 6.6 % | 38,183 | M<sub>200</sub>V | Unknown |
| Shieh | *b* | 19.4 % | 24,026 | V<sub>300</sub>I | Unknown* |
| Shieh | *b* | 6.9 % | 40,807 | K<sub>10</sub>K | DNA helicase |

Bacterial genomes were investigated on the isolate (clonal) level. Isolates were picked from different treatments at several time points during the experiment. Evidence of surface modification was found in almost all phage-exposed samples in the flavobacterial gliding motility genes which are associated with type IX secretion system and whose mutations are expected to cause colony morphotype change from Rhizoid to Rough^45^. Most of these mutations caused premature stop codons or introduced frameshift mutations (Table 2). Most of the T9SS mutants were Rough (Table 2). Isolates from the control treatments without phage did not show mutations in gliding motility genes but had variation in other ORFs. It is therefore possible that extended growth in these conditions introduced other adaptive changes, although these mutations were not present in the phage-exposed samples. Indeed, most variation was detected in the Shieh + M control isolate, which seemed to have had developed resistance against phage even in the phage’s absence (Figure 2B). This sample contained non-synonymous mutations in several metabolism related genes (e.g. *Lon, rpoB, surE*), and in a putative type VI secretion system -like gene (Supplementary File 2). Type VI secretion systems have previously been shown to be associated with host colonization and bacterial antagonism in *Flavobacterium johnsoniae*^46^. Mutations in the *gld* and *spr* genes resulted in resistance against the ancestral V156 phage.

**Table 2.** Bacterial mutations in gliding motility related genes. For a detailed list of all mutations, see supplementary File 2. Resistance: R indicates complete phage resistance, and numbers indicate the last phage dilution that caused observable plaques in the bacterial lawn (phage V156 stock concentration 10^10^ pfu mL^-1^).

| Treatment (replicate) | Week | Morphotype | Gliding motility mutation | Mutation type | Resistance |
| --- | --- | --- | --- | --- | --- |
| LW (b) | 4 | Rough | sprA | Stop codon | R |
| LW (b) | 4 | Rough | sprA | Stop codon | R |
| LW (b) | 8 | Rough | sprA | Stop codon | R |
| LW (c) | 15 | Rough | gldM | Stop codon | R |
| LW+M (a) | 15 | Rough | gldK | SNP nonsynonymous | R |
| LW+M (b) | 12 | Rhizoid | gldK (intergenic) | Intergenic poly-AT | R |
| LW+M (b) | 15 | Rhizoid | sprA | Single nt deletion -> frameshift middle of gene | R |
| LW+M (c) | 8 | Rhizoid | *none* | | $10^{-2}$ |
| Shieh (a) | 12 | Rough | sprA, SusC/RagA TonB-linked outer membrane protein | sprA: poly-A 8->7. SusC: 48bp deletion | R |
| Shieh (b) | 8 | Rough | *none* |  | R |
| Shieh (c) | 10 | Rhizoid | *none* | | $10^{-3}$ |
| Shieh+M (a) | 12 | Rhizoid | ompA | SNP nonsynonymous | $10^{-3}$ |
| Shieh+M (a) | 8 | Rough | ompA, sprE | ompA: (CCATCA)repeat 3->2. sprE: poly-A 7->8 | R |
| Shieh+M (c) | 8 | Rough | GldN, gliding motility related gene (LOCUS_17550) | GldN: 1bp deletion -> frameshift. LOCUS 17550: 42bp deletion | R |
| LW (ctrl) | 3 | Rhizoid | *none* | | $10^{-2}$ |
| LW+M (ctrl) | 15 | Rough | *none* | | $10^{-5}$ |
| Shieh (ctrl) | 12 | Rough | *none* | | $10^{-5}$ |
| Shieh+M (ctrl) | 15 | Rhizoid | Putative T6SS -related gene | | $10^{-5}$ |
| Ancestor B245 | | Rhizoid | | | $10^{-5}$ |

## Discussion

Environmentally transmitted opportunistic bacteria survive long periods in the environment, which may select for reduced growth and metabolic rate^47^. Whereas the interactions between pathogenic bacteria and their phages are generally studied in well-defined laboratory conditions, in real life they often interact in the complex mucosal surfaces of the vertebrate host. Despite the central role of mucosal surfaces for health, there are several fundamental gaps in our knowledge regarding the biology of the phage-bacterium interactions in the mucosal environment. Chemical signals of the host regulates bacterial genes needed for invasion and virulence^34^, generating a physiological state exploited by mucosal phages for infection^7^. Moreover, the ability to colonize hosts is often associated with trade-offs in phage resistance^22^. It is therefore relevant to ask how mucosal environment affects relative investment into different bacterial defense mechanisms against phages. Here, we investigated this question using the opportunistic pathogen *F. columnare* that causes mucosal disease in freshwater fish^30^ but is known to be resilient and able to withstand adverse conditions outside the host^47-48^. Our study suggests that mucin has a central role in triggering CRISPR-Cas immunity and in buffering bacterial survival.

The lake water conditions with and without mucin (LW+M, LW) used in this study are the closest approximations of natural conditions for *F. columnare* thus far. The starved bacterial population (LW) relied almost completely on extracellular immunity via surface modifications, leading to steady decline in the bacterial population density over the 16-week experiment. As the control isolate went extinct in the LW treatment but phage-exposed replicates did not, it is possible that phages have a positive effect for the long term survival of the bacteria population during starvation, perhaps through resistant cells feeding on lysed ones or through Cas13-induced dormancy by the VI-B CRISPR-Cas locus^49^. However, this result could be a fluctuation in our experiment due to having just one control culture and requires further investigation. Presence of mucin (LW+M), on the other hand, triggered the intracellular CRISPR-Cas systems (as measured by spacer acquisition), and supported higher bacterial population densities. The extent of spacer acquisition in this condition was also positively affected by the presence of a competitor bacterium, highlighting that the surrounding microbiome may also play a synergistic role with mucin in determining defense strategies. Presence of mucin also led to more dramatic decline in phage titers, possibly through active removal of phages from the environment using the CRISPR-Cas systems^50^. In rich medium (Shieh), mucin also accelerated spacer acquisition during the first half of the experiment but with much lesser effect, while medium without mucin did not lead to spacer acquisition at all.

These results suggest that the role of CRISPR-Cas in *F. columnare* may be important specifically during colonization of the metazoan. In this setting, the fitness loss associated with surface modification is amplified by reduced bacterial virulence, as colony morphotype change leads to loss of virulence^33,51-52^ via mutations in the flavobacterial gliding motility genes associated with type 9 secretion system^45^. Similarly to previous findings^53^, most phage-exposed bacterial isolates collected in this experiment had mutations in *gld* or *spr* genes involved in the secretion of adhesins on the cell surface. These mutations also occurred in the LW+M treatment, indicating that CRISPR-Cas is not the sole resistance mechanism even in this condition. However, the phage concentration in our experimental system was likely higher than on natural fish surfaces, in which CRISPR-Cas mediated defense may be enough to carry the bacterium through the colonization process without compromising virulence through SM.

In general, bacteria isolated from all treatments throughout the 16-week experiment showed increased resistance to phage. Resistance was especially associated with the Rough colony morphotype. Surprisingly, the experimental conditions had an impact on the benefits of acquired CRISPR spacers.

Expanded CRISPR arrays had a positive effect on bacterial growth in Rhizoid colonies (but negative effects on Rough colonies) in isolates originating from the LW+M treatment (Figure 5A). However, in isolates from Shieh+M treatment, the result was opposite: Rough isolates had higher population density than Rhizoid with and without new spacers. Why specifically rhizoid LW+M samples gained the most benefit from new spacers is unclear, but may reflect an altered bacterial metabolic state induced by the mucin signals indicating the presence of the fish host. If this phenomenon is widespread in bacterial pathogens, the finding may have important real-life implications for the phage-bacterium interactions in metazoan hosts, which are not observed in standard laboratory cultures. Together, these results show that the benefit from additional spacers is affected by the environment where the acquisition happened, and that spacers may even have a negative effect depending on simultaneous surface modification. It is, however, worth noting that the growth experiment was performed in the nutrient-rich Shieh medium, which may downplay the activity of CRISPR-Cas as shown in the 16-week experiment.

Previous studies indicate that high diversity of CRISPR spacers can drive phage populations into extinction^54^. However, our phage-bacterium system behaved differently. Despite the selection pressure by surface modifications and CRISPR-Cas defense, phage populations showed remarkably high genetic stability through the entire experiment, as clearly mutated phage genomes were found only in two replicates. In both cases, genetic change was observed in structural genes, indicating selection imposed by surface modification-based resistance, as seen also previously^55^. Both SM and CRISPR-driven changes in phage genomes have been observed also in the environment^31^, but despite the high prevalence of new CRISPR spacers in the LW+M treatment, no phage mutations were found in this treatment. We also did not find mutations in the pre-existing B245 spacer targets in any treatment. The lack of CRISPR evasion points to either inefficient interference, or an efficient degradation of phages with no leakage of mutated phages. The evolutionary potential of these phages may also be limited, as suggested by slow emergence of mutations in natural samples over several years in this phage-bacterium system^31^.

Diversity in bacterial communities has been shown to increase the benefits of CRISPR immunity in *P. aeruginosa*^29^. We obtained similar results with *F. columnare*, although our study focused only in CRISPR adaptation. Co-culture with another aquatic bacterium, *Aeromonas sp*., led to higher number of spacers acquired, especially in the Rhizoid colony type, indicating a possible trade-off between surface resistance and competitive ability.

Our results support the view that phage defense strategies are influenced by ecological determinants, similar to previous studies showing the effects of nutrient and phage concentrations or the presence of competing species^1,3,29^. In our study system, the bacterium increased intracellular defenses in the presence of metazoan host-signals, which may be an adaptive maneuver to maximize colonization by avoiding the predation by mucosal-associated phages. The necessity to avoid surface modification in mucosal settings may select for diversification of immune mechanisms, and partly explain why some CRISPR-Cas systems are enriched in pathogens^24^. It will be interesting to see if these results are generalizable to other phage-bacterium-metazoan systems. Furthermore, the roles of other intracellular systems besides CRISPR-Cas, such as restriction modification, should be investigated. From a practical viewpoint, understanding the interplay of bacterial virulence and phage defense during bacterial colonization of mucosal surfaces is crucial for the development of phage therapy, which specifically functions in this tripartite scenario.

## Acknowledgements

We would like to thank MSc Katie Smith and Mr Petri Papponen for help in the laboratory, and Dr. Elina Laanto for donating bacterial and phage isolates. This study was funded by the Academy of Finland grant #314939.

## Notes

### Competing Interest Statement

The authors have declared no competing interest.

