## supplementary text, supplementary figures 1-3 for "Mucin induces CRISPR-Cas defence in an opportunistic pathogen"

This file contains Supplementary text and Supplementary Figures 1-3.

**Supplementary text**

***Statistics/Data analysis***

*Effect of treatment of phage resistance*

R library glmmTMB^1^ was used to analyze how the isolates differ from the ancestral bacterium in the presence or absence of phage. We used maximum OD reached during the follow-up growth experiment (OD_MAX_) as the response variable and the interaction between phage and treatment as the explanatory variables. We included replicate and bacterial ID as random effects and used a gaussian distribution as the response variable was normally distributed. The control samples were investigated using a linear model (lm in base-R) using the same interaction terms (the same random structure as in the phage-exposed samples could not be used due to limited sample size in the control dataset).

*Differences in titers of LW and LW+M*

To reveal the effect of mucin on growth in lake water, we analyzed phage and bacterial titers from these conditions using a linear model in R. Titer was set as the response variable and treatment (LW or LW+M) as the explanatory variable. For phage titers, we only considered time points after 9 weeks, as the divergence between the two treatments started at this time (Supplementary Figure 2A). As the bacterial titers diverged starting from week 1 (Supplementary Figure 2B), we included all time points in the model. Similar analysis was not done for the Shieh and Shieh + mucin due to large variation of replicate cultures in these conditions.

*The effect of spacers and morphology*

To analyze the effects of spacers and morphology on OD_MAX_ in the LW+M and Shieh+M treatments, we created a generalized linear mixed model (GLMM) using the R package glmmTMB^1^. Both treatments were studied separately and categorized as having no new spacers or having one or more spacers. OD_MAX_ was used as the response variable and predictors were [the presence of spacers] * [morphology]. As random effects we included plate ID (the plate on which each sample was measured on) and replicate, which both increased model-fit considerably. A gamma distribution of the response variable was favored over other distributions with AIC difference > 2.

*The effect of phage on control samples*

We examined if OD_MAX_ of the control samples (grown in the four treatments without phage) in the growth experiment behaved similarly to the ancestral bacterium. We used a linear model (lm) in R with OD_MAX_ as the response variable and treatment as the explanatory variable. The ancestral bacterium was used as intercept against which all treatments were compared to.

*Effect of extraction time on OD_MAX_*

Using a linear mixed model, we investigated the effect of extraction time on OD_MAX_ and OD_T_. Including replicate as a random factor produced the best models according to AIC. None of the samples showed a significant effect of isolation time on either response variable in the presence or absence of phage (Supplementary Figure 3).

*Effect of competition on F. columnare spacer acquisition*

For the statistical analysis of the competition experiment, we used a generalized linear model (glmmTMB package) with total number of new spacer per colony as the response variable and condition (co-culture or not) as the explanatory variable. Replicate ID and sampling day were used as random effects. AIC model comparison of several distributions (including zero inflated ones) indicated that a negative binomial distribution (nbinom2) fit the data best.

**Supplementary figures**

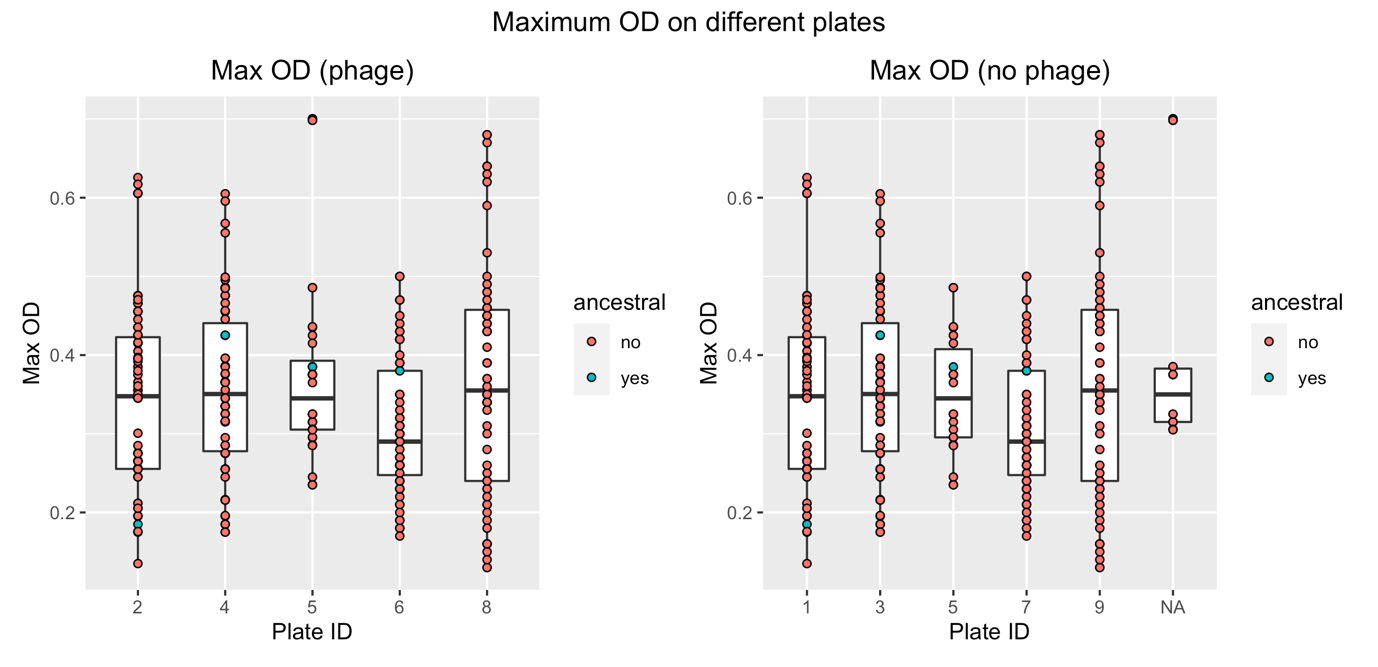

**Supplementary Figure 1.** Effects of measurement plates on growth in Bioscreen.

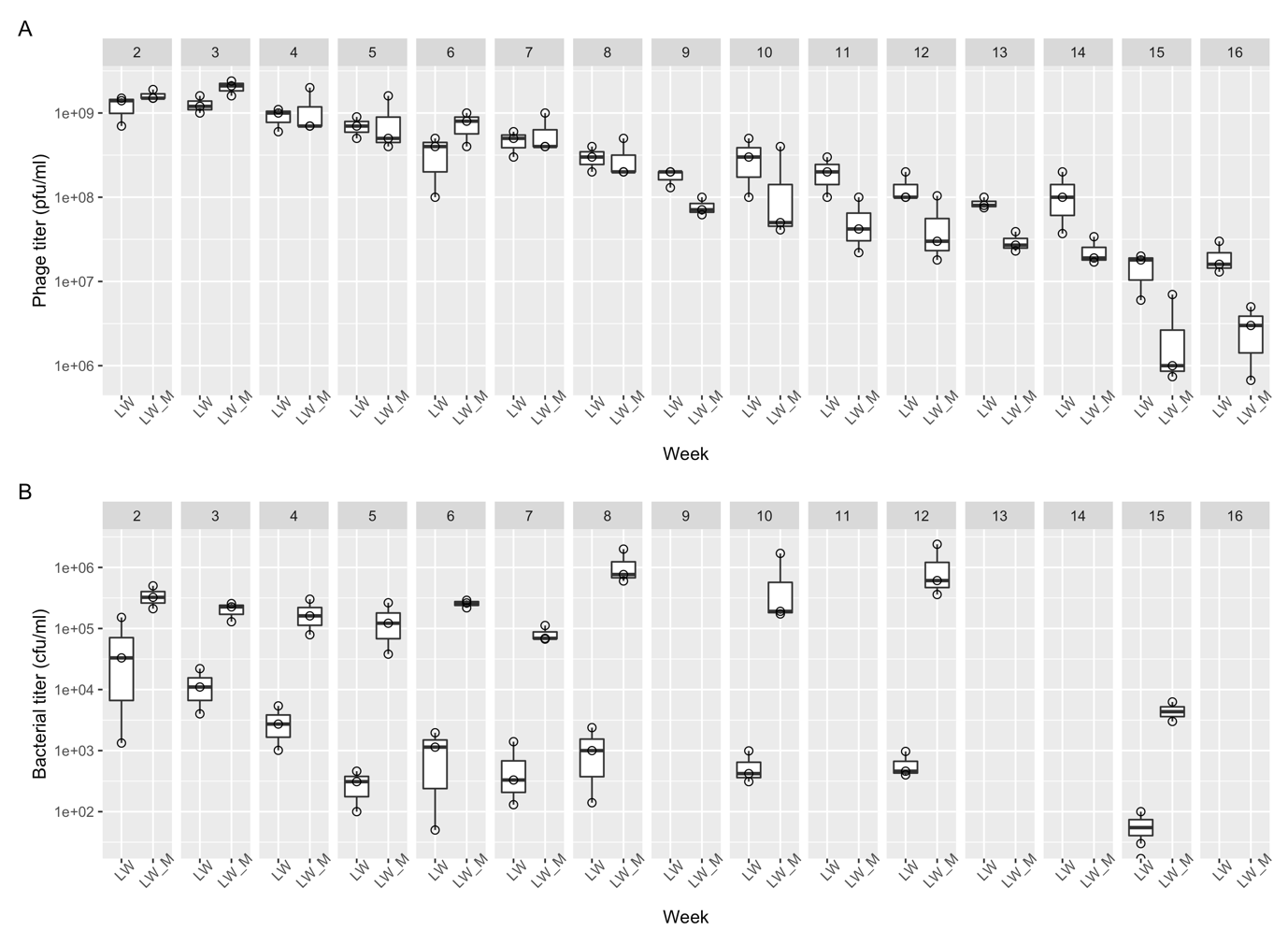

**Supplementary Figure 2.** Comparison of LW and LW+M phage and bacterial titers

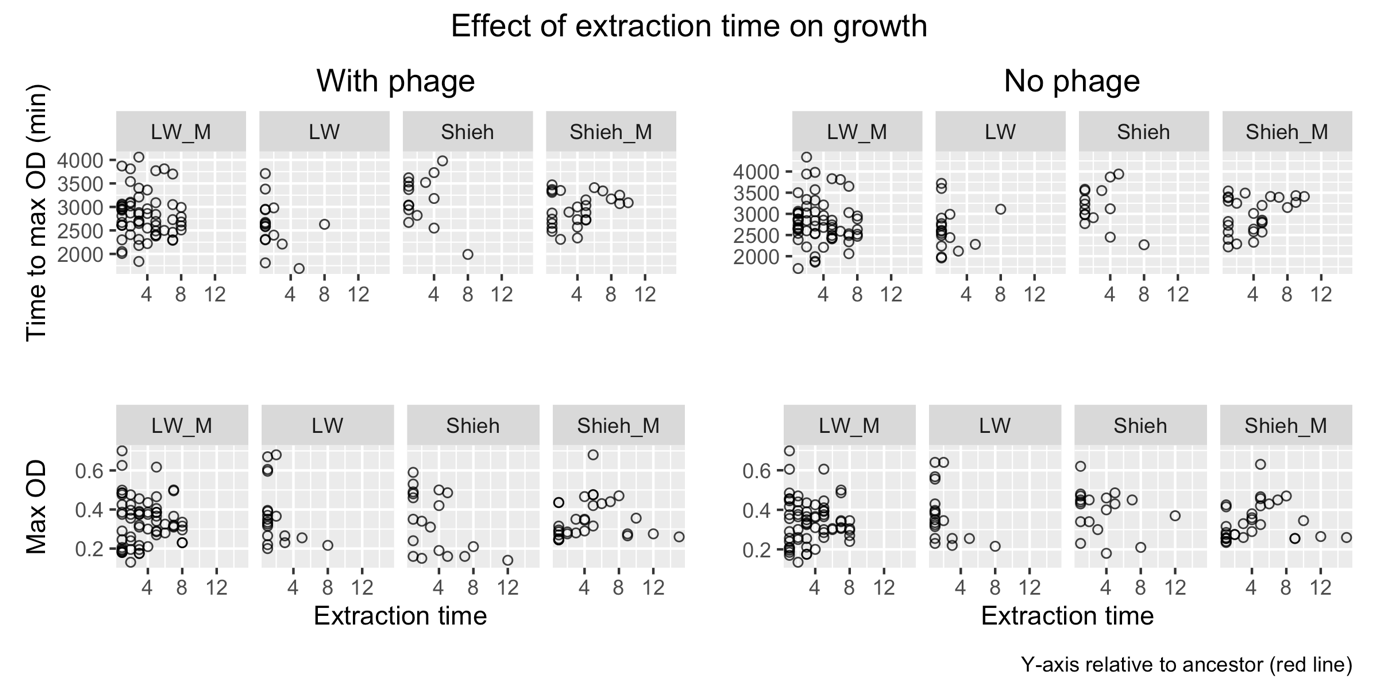

**Supplementary Figure 3.** The effect of extraction time on OD-max and time-to-OD-max.

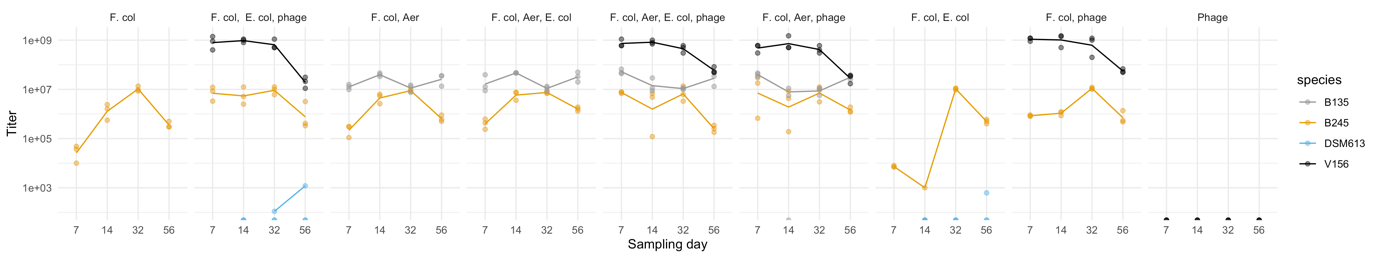

**Supplementary Figure 4.** Bacterial and phage titers from the competition experiments
