## Supplementary file 1 for "Mucin induces CRISPR-Cas defence in an opportunistic pathogen": Supplementary File 1.html

Mutation Comparison


| Predicted mutations | | | | | | | | | | | | | | | | | |
| --- | --- | --- | --- | --- | --- | --- | --- | --- | --- | --- | --- | --- | --- | --- | --- | --- | --- |
| position | mutation | A | B | C | D | F | G | H | I | J | K | L | v156 | fB245 | annotation | gene | description |
| 22,215 | G→A |  |  |  |  |  |  |  |  |  |  | 11.3% |  |  | A357T (GCT→ACT) | *unknown* → | hypothetical protein |
| 24,026 | G→A |  |  |  |  |  |  | 19.4% |  |  |  |  |  |  | V300I (GTA→ATA) | *unknown* → | hypothetical protein |
| 28,227 | C→A | 8.5% | 10.4% | 9.1% | 9.3% | 8.6% | 7.8% | 26.0% | 8.3% | 12.9% | 8.6% | 18.7% |  |  | S340Y (TCT→TAT) | *unknown* → | hypothetical protein |
| 37,372 | G→C | 8.1% | 8.6% | 8.0% | 8.5% | 8.3% |  | 7.1% | 8.4% | 8.6% |  | 7.2% |  | 100% | V171L (GTA→CTA) | *unknown* → | hypothetical protein |
| 38,779 | G→A |  | 7.9% |  |  |  |  |  |  |  |  | 6.7% |  | 100% | L188L (TTG→TTA) | *unknown* → | hypothetical protein |
| 38,797 | T→A |  |  |  |  |  |  |  |  |  |  |  |  | 100% | I194I (ATT→ATA) | *unknown* → | hypothetical protein |
| 38,809 | T→C |  |  |  |  |  |  |  |  |  |  |  |  | 100% | Y198Y (TAT→TAC) | *unknown* → | hypothetical protein |
| 38,813 | A→G |  |  |  |  |  |  |  |  |  |  | 6.6% |  | 100% | M200V (ATG→GTG) ‡ | *unknown* → | hypothetical protein |
| 38,815 | G→T | 7.2% |  | 6.8% | 7.2% | 7.2% | 6.7% | 5.8% | 6.8% | 7.2% | 7.2% | 6.4% |  | 100% | M200I (ATG→ATT) ‡ | *unknown* → | hypothetical protein |
| 38,827 | G→A |  |  |  |  |  |  |  |  |  |  |  |  | 100% | T204T (ACG→ACA) | *unknown* → | hypothetical protein |
| position | mutation | A | B | C | D | F | G | H | I | J | K | L | v156 | fB245 | annotation | gene | description |
| 38,851 | T→A |  |  |  |  |  |  |  |  |  |  |  |  | 100% | G212G (GGT→GGA) | *unknown* → | hypothetical protein |
| 38,875 | G→A | 7.6% | 7.7% | 7.2% | 7.4% |  | 7.0% | 6.2% | 7.4% | 7.4% |  | 7.0% |  | 100% | E220E (GAG→GAA) | *unknown* → | hypothetical protein |
| 38,884 | C→T | 7.4% | 7.4% | 7.0% | 7.2% |  | 6.7% | 6.1% | 7.2% | 7.2% |  | 6.7% |  | 100% | V223V (GTC→GTT) | *unknown* → | hypothetical protein |
| 38,887 | T→A |  |  |  |  |  |  |  |  |  |  |  |  | 100% | P224P (CCT→CCA) | *unknown* → | hypothetical protein |
| 38,902 | G→A |  | 7.8% |  |  |  |  |  |  |  |  | 6.9% |  | 100% | L229L (TTG→TTA) | *unknown* → | hypothetical protein |
| 38,932 | G→T |  | 8.2% | 7.3% | 7.8% |  | 7.4% | 6.5% |  |  |  | 7.5% |  | 100% | G239G (GGG→GGT) | *unknown* → | hypothetical protein |
| 38,938 | T→C |  |  |  |  |  |  |  |  |  |  | 7.6% |  | 100% | N241N (AAT→AAC) | *unknown* → | hypothetical protein |
| 39,124 | T→C |  |  |  |  |  |  |  |  |  |  |  |  | 100% | S303S (TCT→TCC) | *unknown* → | hypothetical protein |
| 39,248 | C→A | 7.1% |  | 7.7% | 7.8% | 8.3% | 7.6% | 6.5% | 7.8% | 8.2% | 7.8% | 7.2% |  | 100% | S23Y (TCT→TAT) | *unknown* → | hypothetical protein |
| 39,255 | T→C |  |  |  | 7.7% |  |  |  |  |  |  |  |  | 100% | Y25Y (TAT→TAC) | *unknown* → | hypothetical protein |
| position | mutation | A | B | C | D | F | G | H | I | J | K | L | v156 | fB245 | annotation | gene | description |
| 39,276 | T→C |  |  |  | 7.7% |  |  |  |  |  |  | 7.0% |  | 100% | N32N (AAT→AAC) | *unknown* → | hypothetical protein |
| 39,462 | T→C |  |  |  |  |  |  |  |  |  |  |  |  | 100% | A94A (GCT→GCC) | *unknown* → | hypothetical protein |
| 39,485 | T→G |  |  |  |  |  |  |  |  |  |  |  |  | 100% | I102S (ATC→AGC) | *unknown* → | hypothetical protein |
| 39,659 | A→G |  |  |  |  |  |  |  |  |  |  |  |  | 100% | I21V (ATT→GTT) | *unknown* → | hypothetical protein |
| 39,749 | G→A |  |  |  | 7.3% |  |  |  |  | 6.9% |  |  |  | 100% | E51K (GAA→AAA) | *unknown* → | hypothetical protein |
| 39,775 | T→A |  |  |  | 5.3% |  |  |  |  |  |  |  |  | 100% | T59T (ACT→ACA) | *unknown* → | hypothetical protein |
| 39,779 | T→C |  |  |  | 5.4% |  |  |  |  |  |  |  |  | 100% | L61L (TTA→CTA) ‡ | *unknown* → | hypothetical protein |
| 39,781 | A→G |  |  |  | 5.4% |  |  |  |  |  |  |  |  | 100% | L61L (TTA→TTG) ‡ | *unknown* → | hypothetical protein |
| 39,783 | T→A |  |  |  | 5.4% |  |  |  |  | 5.3% |  |  |  | 100% | F62Y (TTT→TAT) | *unknown* → | hypothetical protein |
| 40,749 | G→A |  |  |  |  | 5.2% |  |  |  |  | 5.1% |  |  |  | intergenic (+57/‑29) | *unknown* → / → *unknown* | hypothetical protein/hypothetical protein |
| position | mutation | A | B | C | D | F | G | H | I | J | K | L | v156 | fB245 | annotation | gene | description |
| 40,807 | A→G |  |  |  |  |  |  | 6.9% |  |  |  |  |  | 100% | K10K (AAA→AAG) | *unknown* → | hypothetical protein |
| 40,838 | T→G |  |  |  | 8.4% |  |  |  |  |  |  |  |  | 100% | Y21D (TAT→GAT) | *unknown* → | hypothetical protein |
| 40,843 | C→A |  | 8.4% | 7.8% | 8.1% | 8.3% | 7.7% | 6.8% | 7.7% | 8.4% | 7.6% |  |  |  | G22G (GGC→GGA) | *unknown* → | hypothetical protein |
| 40,843 | 2 bp→AA |  |  |  |  |  |  |  |  |  |  |  |  | 100% | coding (66‑67/525 nt) | *unknown* → | hypothetical protein |
| 40,844 | G→A | 7.6% | 8.3% | 7.8% | 8.0% | 8.2% | 7.6% | 6.7% | 7.7% | 8.3% |  |  |  |  | E23K (GAA→AAA) | *unknown* → | hypothetical protein |
| 40,852 | C→A | 8.3% | 8.9% | 8.5% | 8.6% | 8.9% | 8.2% | 7.1% | 8.3% | 9.0% |  |  |  | 100% | V25V (GTC→GTA) | *unknown* → | hypothetical protein |
| 40,858 | A→T | 8.4% | 9.1% | 8.6% | 8.6% |  | 8.2% | 7.2% | 8.4% |  |  |  |  | 100% | V27V (GTA→GTT) | *unknown* → | hypothetical protein |
| 40,906 | C→A | 8.3% | 9.2% | 8.6% |  |  | 8.3% | 7.3% |  |  |  |  |  | 100% | T43T (ACC→ACA) | *unknown* → | hypothetical protein |
| 40,915 | T→C |  |  |  |  |  |  |  |  |  |  |  |  | 100% | T46T (ACT→ACC) | *unknown* → | hypothetical protein |
| 40,921 | A→C | 8.2% | 9.2% | 8.7% |  |  | 8.3% | 7.3% | 8.6% |  |  |  |  | 100% | I48I (ATA→ATC) | *unknown* → | hypothetical protein |
| position | mutation | A | B | C | D | F | G | H | I | J | K | L | v156 | fB245 | annotation | gene | description |
| 40,951 | A→T | 8.2% | 9.2% | 8.6% |  |  | 8.2% | 7.1% | 8.4% | 9.0% |  |  |  | 100% | A58A (GCA→GCT) | *unknown* → | hypothetical protein |
| 40,963 | A→G |  |  |  |  |  |  |  |  |  |  |  |  | 100% | Q62Q (CAA→CAG) | *unknown* → | hypothetical protein |
| 40,969 | G→A | 8.4% |  | 8.8% | 9.0% |  | 8.5% |  |  | 9.3% |  |  |  | 100% | R64R (AGG→AGA) | *unknown* → | hypothetical protein |
| 40,981 | G→A | 8.3% | 9.2% | 8.7% | 8.9% |  | 8.4% |  | 8.7% | 9.1% |  |  |  | 100% | L68L (TTG→TTA) | *unknown* → | hypothetical protein |
| 41,023 | G→A | 8.1% | 9.0% | 8.4% | 8.9% | 9.2% | 8.3% | 7.2% | 8.6% | 9.0% |  | 7.8% |  | 100% | L82L (TTG→TTA) | *unknown* → | hypothetical protein |
| 41,038 | A→G |  |  |  |  |  |  |  |  |  |  |  |  | 100% | T87T (ACA→ACG) | *unknown* → | hypothetical protein |
| 41,041 | T→A |  |  |  |  |  |  |  |  |  |  |  |  | 100% | V88V (GTT→GTA) | *unknown* → | hypothetical protein |
| 41,053 | T→C |  |  |  |  |  |  |  |  |  |  |  |  | 100% | D92D (GAT→GAC) | *unknown* → | hypothetical protein |
| 41,063 | A→G |  |  |  |  |  |  |  | 8.8% |  | 8.9% | 8.1% |  | 100% | I96V (ATT→GTT) | *unknown* → | hypothetical protein |
| 41,068 | T→A | 8.5% | 9.1% | 8.5% | 9.1% | 9.6% | 8.3% | 7.3% | 8.9% | 9.2% | 9.0% | 8.1% |  | 100% | I97I (ATT→ATA) | *unknown* → | hypothetical protein |
| position | mutation | A | B | C | D | F | G | H | I | J | K | L | v156 | fB245 | annotation | gene | description |
| 41,087 | T→C |  |  |  |  |  |  |  |  |  |  |  |  | 100% | L104L (TTG→CTG) | *unknown* → | hypothetical protein |
| 41,246 | G→A | 8.1% |  | 8.0% | 8.6% |  | 7.8% | 6.7% | 8.2% | 8.5% | 8.5% | 8.0% |  | 100% | A157T (GCT→ACT) | *unknown* → | hypothetical protein |
| 41,259 | G→T | 7.9% |  | 7.8% | 8.4% |  | 7.6% | 6.5% | 7.9% | 8.3% | 8.2% | 7.8% |  | 100% | S161I (AGC→ATC) | *unknown* → | hypothetical protein |
| 41,261 | A→G |  |  |  |  |  |  |  |  |  |  |  |  | 100% | R162G (AGA→GGA) | *unknown* → | hypothetical protein |
| 41,266 | G→A | 8.1% |  | 8.1% | 8.8% |  |  | 6.7% | 8.1% |  | 8.4% | 8.2% |  | 100% | L163L (TTG→TTA) | *unknown* → | hypothetical protein |
| 41,284 | A→C |  | 8.6% |  |  |  |  |  |  |  |  | 8.2% |  | 100% | K169N (AAA→AAC) | *unknown* → | hypothetical protein |
| 41,287 | G→A | 8.1% | 8.5% | 7.9% | 8.8% |  | 7.5% | 6.7% | 8.1% | 8.6% | 8.4% | 8.1% |  | 100% | L170L (TTG→TTA) | *unknown* → | hypothetical protein |
| 41,398 | C→T | 7.7% | 8.2% | 7.8% | 7.9% | 7.8% | 7.5% | 6.4% | 8.0% | 8.3% | 7.9% | 7.5% |  | 100% | L14F (CTT→TTT) | *unknown* → | hypothetical protein |
| 41,480 | G→A | 7.7% | 8.4% | 7.9% | 8.0% | 7.7% | 7.8% | 6.5% | 8.0% | 8.3% | 7.9% | 7.5% |  | 100% | G41D (GGT→GAT) | *unknown* → | hypothetical protein |
| 41,488 | C→T | 7.6% | 8.4% | 7.9% | 8.0% | 7.6% | 7.8% | 6.5% | 7.9% | 8.2% | 8.0% | 7.4% |  | 100% | H44Y (CAT→TAT) | *unknown* → | hypothetical protein |
| position | mutation | A | B | C | D | F | G | H | I | J | K | L | v156 | fB245 | annotation | gene | description |
| 41,491 | G→A | 7.4% | 8.2% | 7.8% |  | 7.5% | 7.6% | 6.4% | 7.8% | 8.0% | 7.8% |  |  | 100% | E45K (GAA→AAA) | *unknown* → | hypothetical protein |
| 41,514 | G→A | 7.7% |  | 8.2% | 7.9% | 7.9% | 7.8% | 6.7% | 8.0% | 8.2% | 8.1% | 7.5% |  | 100% | R52R (AGG→AGA) | *unknown* → | hypothetical protein |
| 42,225 | T→G | 94.3% | 94.6% | 100% | 100% | 100% | 100% | 78.7% | 100% | 93.4% | 100% | 84.4% | 26.6% | 100% | D15E (GAT→GAG) | *unknown* → | hypothetical protein |
| 42,250 | G→A | 100% | 100% | 100% | 100% | 100% | 100% | 100% | 100% | 100% | 100% | 100% |  | 100% | V24I (GTA→ATA) | *unknown* → | hypothetical protein |
| 42,517 | +CGCAACCACAGCAACAAC |  |  |  |  |  |  |  |  |  |  |  |  | 7.5% | coding (337/429 nt) | *unknown* → | hypothetical protein |
| 42,519 | +CAACCACAGCAACAAACT |  |  |  |  |  |  |  |  |  |  |  |  | 28.9% | coding (339/429 nt) | *unknown* → | hypothetical protein |
| 42,519 | +CAACCACAGCAACAACAT |  |  |  |  |  |  |  |  |  |  |  |  | 33.7% | coding (339/429 nt) | *unknown* → | hypothetical protein |
| 42,519 | 17 bp→35 bp |  |  |  |  |  |  |  |  |  |  |  |  | 70.9% | coding (339‑355/429 nt) | *unknown* → | hypothetical protein |
| 42,519 | 17 bp→35 bp |  |  |  |  |  |  |  |  |  |  |  |  | 55.9% | coding (339‑355/429 nt) | *unknown* → | hypothetical protein |
| 42,519 | 19 bp→37 bp |  |  |  |  |  |  |  |  |  |  |  |  | 66.9% | coding (339‑357/429 nt) | *unknown* → | hypothetical protein |
| position | mutation | A | B | C | D | F | G | H | I | J | K | L | v156 | fB245 | annotation | gene | description |
| 42,519 | 19 bp→37 bp |  |  |  |  |  |  |  |  |  |  |  |  | 42.1% | coding (339‑357/429 nt) | *unknown* → | hypothetical protein |
| 42,519 | 28 bp→46 bp |  |  |  |  |  |  |  |  |  |  |  |  | 64.8% | coding (339‑366/429 nt) | *unknown* → | hypothetical protein |
| 42,519 | 29 bp→46 bp |  |  |  |  |  |  |  |  |  |  |  |  | 13.1% | coding (339‑367/429 nt) | *unknown* → | hypothetical protein |
| 42,520 | +AACCACAGCAACAACCTA |  |  |  |  |  |  |  |  |  |  |  |  | 33.7% | coding (340/429 nt) | *unknown* → | hypothetical protein |
| 42,543 | +CCTCAACAAGCACAACAA |  | 5.3% |  |  | 6.8% |  |  | 6.5% |  |  |  |  |  | coding (363/429 nt) | *unknown* → | hypothetical protein |
| 42,662 | C→T | 100% | 100% | 100% | 100% | 100% | 100% | 100% | 100% | 93.6% | 100% | 100% |  | 100% | intergenic (+53/‑69) | *unknown* → / → *unknown* | hypothetical protein/hypothetical protein |
| 42,699 | A→T | 94.5% | 94.7% | 94.9% |  |  |  |  |  |  |  | 84.3% |  |  | intergenic (+90/‑32) | *unknown* → / → *unknown* | hypothetical protein/hypothetical protein |
| 42,699 | 2 bp→TA |  |  |  | 100% | 100% | 100% | 100% | 100% | 100% | 100% |  |  | 100% | intergenic (+90/‑31) | *unknown* → / → *unknown* | hypothetical protein/hypothetical protein |
| 42,700 | G→A | 94.5% | 94.8% | 95.0% |  |  |  |  |  |  |  | 100% |  |  | intergenic (+91/‑31) | *unknown* → / → *unknown* | hypothetical protein/hypothetical protein |
| 42,703 | G→A | 94.6% | 94.9% | 100% | 100% | 100% | 100% | 100% | 100% | 100% | 100% | 100% |  | 100% | intergenic (+94/‑28) | *unknown* → / → *unknown* | hypothetical protein/hypothetical protein |
| position | mutation | A | B | C | D | F | G | H | I | J | K | L | v156 | fB245 | annotation | gene | description |
| 42,706 | G→C | 94.6% | 94.9% | 100% | 100% | 100% | 100% | 100% | 100% | 100% | 100% | 84.8% |  | 100% | intergenic (+97/‑25) | *unknown* → / → *unknown* | hypothetical protein/hypothetical protein |
| 42,971 | A→G | 100% | 100% | 100% | 100% | 100% | 100% | 100% | 100% | 100% | 100% | 100% |  | 100% | intergenic (+31/‑275) | *unknown* → / → *unknown* | hypothetical protein/hypothetical protein |
| 43,047 | A→T | 94.1% | 100% | 100% | 100% | 100% | 100% | 100% | 100% | 100% | 100% | 84.4% | 26.6% | 100% | intergenic (+107/‑199) | *unknown* → / → *unknown* | hypothetical protein/hypothetical protein |
| 43,098 | C→T | 100% | 100% | 100% | 100% | 100% | 100% | 100% | 100% | 100% | 100% | 100% |  | 100% | intergenic (+158/‑148) | *unknown* → / → *unknown* | hypothetical protein/hypothetical protein |
| 43,265 | C→T |  |  |  |  |  |  |  |  |  |  |  |  | 100% | T7I (ACT→ATT) | *unknown* → | hypothetical protein |
| 48,229 | C→T |  |  |  |  |  |  |  |  |  |  |  |  | 100% | intergenic (+31/–) | *unknown* → / *–* | hypothetical protein/– |
| 48,540 | T→G | 100% | 100% | 100% | 86.8% | 100% | 100% | 72.4% | 88.9% | 82.7% | 100% | 76.7% |  | 100% | intergenic (+342/–) | *unknown* → / *–* | hypothetical protein/– |
