## Supplementary file 2 for "Mucin induces CRISPR-Cas defence in an opportunistic pathogen": Supplementary File 2.html

Mutation Comparison


| Predicted mutations | | | | | | | | | | | | | | | | | | | | | | |
| --- | --- | --- | --- | --- | --- | --- | --- | --- | --- | --- | --- | --- | --- | --- | --- | --- | --- | --- | --- | --- | --- | --- |
| position | mutation | L8-4 | J8-6 | J12-1 | I10-1 | H8-8 | G12-5 | F8-1 | E15-1 | E12-2 | D15-8 | C15-1 | B8-1 | B4-3 | B4-1 | Shctr | ShMctr | LWctr | LWMctr | annotation | gene | description |
| 95,926 | Δ5 bp |  |  |  |  |  |  |  | 100% |  |  |  |  |  |  |  |  |  |  | coding (1204‑1208/2451 nt) | *lon* ← | Lon protease |
| 96,004 | A→G |  |  |  |  |  |  |  |  |  |  |  |  |  |  |  | 100% |  |  | L377S (TTA→TCA) | *lon* ← | Lon protease |
| 163,549 | C→T |  |  |  |  |  |  |  |  |  |  | 100% |  |  |  |  |  |  |  | G162R (GGA→AGA) | *fabH2* ← | 3‑oxoacyl‑[acyl‑carrier‑protein] synthase 3 |
| 163,755 | Δ2 bp |  |  |  |  |  |  |  |  |  |  |  | 100% |  |  |  |  |  |  | coding (277‑278/1008 nt) | *fabH2* ← | 3‑oxoacyl‑[acyl‑carrier‑protein] synthase 3 |
| 490,706 | 101 bp→7 bp | ? | ? | ? | ? | ? | ? | ? | ? | ? | ? | ? |  | ? | ? | ? | ? | 100% | ? | intergenic (‑56/‑495) | *LOCUS\_04380* ← / → *LOCUS\_04390* | hypothetical protein/hypothetical protein |
| 490,707 | 100 bp→6 bp | ? | ? | ? | ? | ? | ? | ? | ? | ? | ? | ? | 100% | ? | ? | ? | ? | ? | ? | intergenic (‑57/‑495) | *LOCUS\_04380* ← / → *LOCUS\_04390* | hypothetical protein/hypothetical protein |
| 510,279 | C→T |  |  |  |  |  |  |  |  |  |  |  |  |  |  |  | 100% |  |  | E319K (GAA→AAA) | *rpoB* ← | DNA‑directed RNA polymerase subunit beta |
| 529,529 | Δ48 bp |  |  |  |  |  | 100% |  |  |  |  |  |  |  |  |  |  |  |  | coding (1877‑1924/2940 nt) | *LOCUS\_04680* ← | SusC/RagA family TonB‑linked outer membrane protein |
| 693,770 | (TTAATAAAACTTCTATC)_1→2_ |  |  |  |  |  |  |  |  | 100% |  |  |  |  |  |  |  |  |  | coding (192/1359 nt) | *LOCUS\_06060* ← | ligase |
| 696,398 | (T)_9→8_ |  |  |  |  |  |  | 100% |  |  |  |  |  |  |  |  |  |  |  | coding (762/1134 nt) | *LOCUS\_06090* ← | glycosyl transferase |
| position | mutation | L8-4 | J8-6 | J12-1 | I10-1 | H8-8 | G12-5 | F8-1 | E15-1 | E12-2 | D15-8 | C15-1 | B8-1 | B4-3 | B4-1 | Shctr | ShMctr | LWctr | LWMctr | annotation | gene | description |
| 727,481 | Δ30 bp |  |  |  |  |  |  |  |  |  |  |  |  |  |  |  |  |  | 100% | coding (83‑112/618 nt) | *LOCUS\_06360* ← | acetyltransferase |
| 799,525 | (A)_10→11_ |  |  |  |  |  |  |  |  |  |  | 100% |  |  |  |  |  |  |  | intergenic (+28/‑97) | *LOCUS\_07050* → / → *LOCUS\_07060* | membrane protein/SusC/RagA family TonB‑linked outer membrane protein |
| 916,603 | Δ1 bp |  |  |  |  |  |  |  | 100% |  |  |  |  |  |  |  |  |  |  | coding (672/1485 nt) | *LOCUS\_07840* ← | hypothetical protein |
| 919,708 | +T | 100% |  |  |  |  |  |  |  |  |  |  |  |  |  |  |  |  |  | coding (24/996 nt) | *gldN* ← | gliding motility protein GldO |
| 920,562 | C→A |  |  |  |  |  |  |  |  |  |  | 100% |  |  |  |  |  |  |  | E228\* (GAA→TAA) | *gldM* ← | gliding motility protein GldM |
| 922,403 | A→G |  |  |  |  |  |  |  |  |  | 100% |  |  |  |  |  |  |  |  | S337P (TCT→CCT) | *gldK* ← | gliding motility lipoprotein GldK |
| 923,503 | (ATTT)_3→2_ |  |  |  |  |  |  |  |  | 100% |  |  |  |  |  |  |  |  |  | intergenic (‑92/+45) | *gldK* ← / ← *fjo29* | gliding motility lipoprotein GldK/arginase |
| 1,103,356 | +T | 100% | 100% | 100% |  |  | 100% | 100% | 100% | 100% | 100% | 100% |  |  |  | 100% |  |  | 100% | coding (86/507 nt) | *LOCUS\_09580* → | thioredoxin |
| 1,108,045 | C→T |  |  |  |  |  |  |  |  |  |  |  |  |  |  |  | 100% |  |  | G13E (GGG→GAG) | *surE* ← | 5'‑nucleotidase SurE |
| 1,112,746 | G→T | ? | ? | ? | ? | ? | ? | ? | ? | ? | ? | ? | ? | ? | ? | ? | ? | 100% | ? | I49I (ATC→ATA) | *LOCUS\_09660* ← | hypothetical protein |
| position | mutation | L8-4 | J8-6 | J12-1 | I10-1 | H8-8 | G12-5 | F8-1 | E15-1 | E12-2 | D15-8 | C15-1 | B8-1 | B4-3 | B4-1 | Shctr | ShMctr | LWctr | LWMctr | annotation | gene | description |
| 1,195,701 | T→C |  |  |  |  |  |  |  |  | 100% |  |  |  |  |  |  |  |  |  | A40A (GCT→GCC) | *LOCUS\_10350* → | hypothetical protein |
| 1,203,521 | Δ1 bp |  |  |  |  |  |  |  | 100% |  |  |  |  |  |  |  |  |  |  | coding (4610/7185 nt) | *sprA* ← | cell surface protein SprA |
| 1,206,903 | C→A |  |  |  |  |  |  |  |  |  |  |  | 100% | 100% | 100% |  |  |  |  | E410\* (GAG→TAG) | *sprA* ← | cell surface protein SprA |
| 1,207,236 | (A)_8→7_ |  |  |  |  |  | 100% |  |  |  |  |  |  |  |  |  |  |  |  | coding (895/7185 nt) | *sprA* ← | cell surface protein SprA |
| 1,301,487 | C→T |  |  |  |  | 100% | 100% |  |  | 100% |  |  | 100% | 100% |  |  |  |  | 100% | C200Y (TGC→TAC) | *LOCUS\_11250* ← | integrase |
| 1,340,764 | C→T | ? | ? | ? |  | ? | ? | ? | ? | ? | ? | ? |  | 100% | ? |  |  | ? | ? | intergenic (‑116/+894) | *LOCUS\_11460* ← / ← *LOCUS\_11470* | hypothetical protein/hypothetical protein |
| 1,344,055 | A→T |  |  |  |  |  |  |  |  |  |  |  |  |  |  |  | 100% |  |  | F326I (TTT→ATT) | *LOCUS\_11480* ← | hypothetical protein |
| 1,418,761 | G→A |  |  |  |  |  |  |  |  |  |  |  |  |  |  |  | 100% |  |  | Q85\* (CAA→TAA) | *LOCUS\_12130* ← | two‑component sensor histidine kinase |
| 1,497,270 | (CCATCA)_3→2_ |  | 100% |  |  |  |  |  |  |  |  |  |  |  |  |  |  |  |  | coding (792‑797/1407 nt) | *LOCUS\_12770* ← | cell envelope biogenesis protein OmpA |
| 1,497,323 | G→A |  |  | 100% |  |  |  |  |  |  |  |  |  |  |  |  |  |  |  | D248Y (GAC→TAT) | *LOCUS\_12770* ← | cell envelope biogenesis protein OmpA |
| position | mutation | L8-4 | J8-6 | J12-1 | I10-1 | H8-8 | G12-5 | F8-1 | E15-1 | E12-2 | D15-8 | C15-1 | B8-1 | B4-3 | B4-1 | Shctr | ShMctr | LWctr | LWMctr | annotation | gene | description |
| 1,497,325 | C→A |  |  | 100% |  |  |  |  |  |  |  |  |  |  |  |  |  |  |  | D248Y (GAC→TAT) | *LOCUS\_12770* ← | cell envelope biogenesis protein OmpA |
| 1,497,332 | T→A |  |  | 100% |  |  |  |  |  |  |  |  |  |  |  |  |  |  |  | Q245H (CAA→CAT) | *LOCUS\_12770* ← | cell envelope biogenesis protein OmpA |
| 1,497,344 | T→A |  |  | 100% |  |  |  |  |  |  |  |  |  |  |  |  |  |  |  | K241N (AAA→AAT) | *LOCUS\_12770* ← | cell envelope biogenesis protein OmpA |
| 1,497,365 | G→A |  |  | 100% |  |  |  |  |  |  |  |  |  |  |  |  |  |  |  | N234N (AAC→AAT) | *LOCUS\_12770* ← | cell envelope biogenesis protein OmpA |
| 1,497,368 | G→T |  |  | 100% |  |  |  |  |  |  |  |  |  |  |  |  |  |  |  | F233L (TTC→TTA) | *LOCUS\_12770* ← | cell envelope biogenesis protein OmpA |
| 1,721,311 | +T |  |  |  |  |  |  |  |  |  | 100% |  |  |  |  |  |  |  |  | coding (221/1341 nt) | *capK* ← | AMP‑binding protein |
| 1,731,445 | (A)_7→8_ |  | 100% |  |  |  |  |  |  |  |  |  |  |  |  |  |  |  |  | coding (876/2607 nt) | *sprE* → | gliding motility protein |
| 1,973,749 | C→T | 100% |  |  |  |  |  |  |  |  |  |  |  |  |  |  |  |  |  | A247V (GCA→GTA) | *ctaD* → | cytochrome‑c oxidase |
| 2,073,773 | Δ42 bp | 100% |  |  |  |  |  |  |  |  |  |  |  |  |  |  |  |  |  | coding (146‑187/207 nt) | *LOCUS\_17550* ← | hypothetical protein |
| 2,132,463 | C→A |  |  |  |  |  |  |  |  |  |  |  |  |  |  |  |  |  | 100% | A108E (GCA→GAA) | *LOCUS\_18070* → | permease |
| position | mutation | L8-4 | J8-6 | J12-1 | I10-1 | H8-8 | G12-5 | F8-1 | E15-1 | E12-2 | D15-8 | C15-1 | B8-1 | B4-3 | B4-1 | Shctr | ShMctr | LWctr | LWMctr | annotation | gene | description |
| 2,134,070 | A→T |  | 100% | 100% |  |  |  | 100% | 100% |  |  |  | 100% |  |  |  |  | 100% | 100% | intergenic (+676/+365) | *LOCUS\_18070* → / ← *LOCUS\_18080* | permease/hypothetical protein |
| 2,246,024 | +66 bp |  |  |  |  |  |  |  |  |  |  |  | 100% |  |  |  |  |  |  | intergenic (‑1607/+137) | *LOCUS\_19110* ← / ← *LOCUS\_19120* | hypothetical protein/hypothetical protein |
| 2,418,824 | C→G |  |  |  | 100% | ? |  |  |  |  |  |  |  |  |  |  | 100% |  |  | G108R (GGC→CGC) | *LOCUS\_20580* ← | Crp/Fnr family transcriptional regulator |
| 2,466,000 | +C | 100% | 100% | 100% | 100% |  |  | 100% | 100% | 100% |  |  | 100% |  |  | 100% | 100% | 100% |  | intergenic (+29/‑51) | *LOCUS\_20950* → / → *LOCUS\_20960* | hypothetical protein/hypothetical protein |
| 2,471,336 | G→A |  |  |  | 100% |  |  |  | 100% |  | 100% |  |  |  |  |  | 100% |  | 100% | D211N (GAT→AAT) | *LOCUS\_21020* → | hypothetical protein |
| 2,471,434 | A→C |  |  | 100% |  |  |  | 100% |  |  |  |  |  |  |  |  | 100% |  |  | T243T (ACA→ACC) | *LOCUS\_21020* → | hypothetical protein |
| 2,472,219 | T→A |  | 100% |  |  |  |  |  |  |  |  |  |  |  |  |  |  |  |  | H74Q (CAT→CAA) | *LOCUS\_21040* → | hypothetical protein |
| 2,472,255 | T→C |  |  |  |  |  |  |  |  |  | 100% |  |  |  |  |  |  |  |  | A86A (GCT→GCC) | *LOCUS\_21040* → | hypothetical protein |
| 2,472,258 | G→A |  | 100% |  |  |  |  |  |  |  | 100% |  |  |  |  |  |  |  |  | E87E (GAG→GAA) | *LOCUS\_21040* → | hypothetical protein |
| 2,557,465 | A→G | 100% | 100% | 100% | 100% | 100% | 100% | 100% | 100% | 100% | 100% | 100% | 100% | 100% | 100% | 100% | 100% | 100% | 100% | \*85W (TGA→TGG) | *LOCUS\_21750* → | hypothetical protein |
| position | mutation | L8-4 | J8-6 | J12-1 | I10-1 | H8-8 | G12-5 | F8-1 | E15-1 | E12-2 | D15-8 | C15-1 | B8-1 | B4-3 | B4-1 | Shctr | ShMctr | LWctr | LWMctr | annotation | gene | description |
| 2,557,473 | A→C | 100% | 100% | 100% | 100% | 100% | 100% | 100% | 100% | 100% | 100% | 100% | 100% | 100% | 100% | 100% | 100% | 100% | 100% | intergenic (+8/‑318) | *LOCUS\_21750* → / → *LOCUS\_21760* | hypothetical protein/hypothetical protein |
| 2,557,529 | Δ1 bp | 100% | 100% | 100% | 100% | 100% | 100% | 100% | 100% | 100% | 100% | 100% | 100% | 100% | 100% | 100% | 100% | 100% | 100% | intergenic (+64/‑262) | *LOCUS\_21750* → / → *LOCUS\_21760* | hypothetical protein/hypothetical protein |
| 2,693,083 | T→G | 100% | 100% | 100% | 100% | 100% | 100% | 100% | ? | 100% | 100% | 100% | 100% | 100% | 100% | 100% | 100% | 100% | 100% | Y40S (TAT→TCT) | *LOCUS\_23110* ← | hypothetical protein |
| 2,693,809 | A→G | 100% | 100% | 100% | 100% |  | ? |  | 100% | 100% |  | 100% |  | 100% | 100% | 100% | 100% | 100% | 100% | I29T (ATA→ACA) | *LOCUS\_23120* ← | transposase |
| 2,747,093 | (T)_8→9_ |  |  |  |  |  | 100% |  |  |  |  |  |  |  |  |  |  |  |  | coding (474/564 nt) | *LOCUS\_23650* ← | DNA polymerase III subunit epsilon |
| 2,903,595 | (T)_5→4_ |  |  | 100% |  |  |  |  |  |  |  |  |  |  |  |  |  |  |  | coding (351/1059 nt) | *LOCUS\_25090* → | hypothetical protein |
| 3,002,261 | G→T |  |  |  |  |  |  |  | 100% |  |  |  |  |  |  |  |  |  |  | N77K (AAC→AAA) | *LOCUS\_25900* ← | hypothetical protein |
